# Kinetic trapping organizes actin filaments within liquid-like protein droplets

**DOI:** 10.1101/2023.05.26.542517

**Authors:** Aravind Chandrasekaran, Kristin Graham, Jeanne C. Stachowiak, Padmini Rangamani

## Abstract

Actin is essential for various cellular functions such as growth, migration, and endocytosis. Recent evidence suggests that several actin-binding proteins phase separate to form condensates and that actin networks have different architectures in these droplets. In this study, we use computational modeling to investigate the conditions under which actin forms different network organizations in VASP droplets. Our simulations reveal that the binding and unbinding rates of actin and VASP determine the probability of formation of shells and rings, with shells being more probable than rings. The different actin networks are highly dependent on the kinetics of VASP-actin interactions, suggesting that they arise from kinetic trapping. Specifically, we showed that reducing the residence time of VASP on actin filaments promotes assembly of shells rather than rings, where rings require a greater degree of actin bundling. These predictions were tested experimentally using a mutant of VASP, which has decreased bundling capability. Experiments reveal an increase in the abundance of shells in VASP droplets, consistent with our predictions. Finally, we investigated the arrangements of filaments within deformed droplets and found that the filament length largely determines whether a droplet will straighten into a bundle or remain kinetically trapped in a ring-like architecture. The sphere-to-ellipsoid transition is favored under a wide range of conditions while the ellipse-to-rod transition is only permitted when filaments have a specific range of lengths. Our findings have implications for understanding how the interactions between phase-separated actin binding proteins and actin filaments can give rise to different actin network architectures.

## Introduction

Phase-separated protein droplets are ubiquitous in cell biology, where they are thought to drive spatial organization within cells by acting as highly localized pockets of signal transduction (1, 2). Phase separation has also been implicated in locally concentrating many proteins that are involved in the remodeling of the actin cytoskeleton (3, 4). Recently, we showed that Vasodilator Stimulated Phosphoprotein (VASP), a processive actin polymerase and an actin bundling protein, forms liquid-like droplets in vitro under physiological conditions (5). When actin was added to these VASP droplets, we found that the actin filaments self-organized into different droplet-encapsulated structures such as shells, rings, or disks. When the actin network formed rings and the ring thickness exceeded a critical value, the actin filament bundle was able to deform the droplet resulting in either an ellipsoidal disk or a linear actin bundle. The energetic arguments in (5) identified the critical ring thickness above which a spherical droplet will deform into an ellipsoid. However, a critical transition prior to the elongation of the droplet is the formation of ring-like structures of actin. When actin was added to these VASP droplets, we found that the actin filaments self-organized into shells, rings, or disks depending on the concentrations of actin and VASP in the droplet. While the conditions under which droplets form have gained significant interest recently (6–8), studies of the downstream kinetic consequences due to phase separation have been limited to transcriptional condensates (8–11). To broaden our understanding of droplet-driven kinetic assembly, here, we study the biophysical mechanisms that give rise to different actin configurations within a VASP droplet.

The interactions of actin with VASP have been studied previously both in cellular (12, 13) and reconstituted systems (14, 15). VASP was originally isolated from platelets (16) where it is known to inhibit platelet activation/aggregation (17, 18). When platelets are exposed to vasodilators, cAMP and cGMP levels are elevated resulting in the kinase-mediated activation of VASP (19, 20). VASP has been subsequently found in several other cell types (21) and has been implicated in cellular motility (22) and axonal guidance (23–25). Subsequent discovery of Ena and Ena-VASP-like (EVL) proteins resulted in a new Ena/VASP family of proteins. The Ena/VASP family of proteins is also of importance clinically due to its role in promoting Listeria motility (26, 27) and cancer cell metastasis (28–30). Ena/VASP proteins are characterized by three major domains: Ena-VASP homology domains (EVH) 1 and 2 separated by a proline rich region. The EVH1 domain contains binding sites for focal adhesion complexes such as Zyxin and Vinculin (27). The EVH2 domain contains the coiled-coil tetramerization domain (31–33), along with G-actin and F-actin binding motifs (34, 35). Therefore, at the molecular level, VASP-tetramers act as weak crosslinkers (36) and processive actin polymerases (14, 37). While the role of Ena/VASP proteins in controlling elongation of actin filaments has been well established, the emergent mechanism of network organization from Ena/VASP crosslinking is not yet fully understood.

Actin filaments form multiple architectures depending on their binding partners and local environment (38, 39). Computational efforts have played a key role in dissecting the biophysics of these processes (40, 41). For example, in cell spreading, actin forms a dendritic branched architecture at the lamellipodium (42) but forms bundled force-generating structures in filopodia (43) and in stress fibers (44, 45). Different actin architectures are also seen in endocytosis (41, 46–48). Each of these network architectures results from a combination of filament-level modifications of various actin binding proteins (49), along with their respective time scales of interaction, which are determined by the kinetics of binding and unbinding (50–54). Our observations with VASP-actin interactions indicate that, depending on the ratio of actin and VASP, different dominant actin architectures can be observed within a VASP droplet. In this work, we sought to investigate the key biophysical determinants of such varied structures using a combination of computational modeling and experiments. Specifically, we hypothesized that different actin networks could be characterized by kinetic accessibility or lack thereof resulting from Actin-VASP interactions.

We used Cytosim, an agent-based modeling framework, (41, 46, 55–57) to model dynamics of the actin filaments confined within an actin droplet at non-equilibrium steady state and investigated how different kinetic time scales including binding and unbinding of actin to VASP, and actin elongation rate affect the formation of actin shells and rings. Our simulations showed that the distribution of rings and shells is determined by the competition between binding and unbinding kinetics of actin and VASP. Other time scales, such as actin elongation rate, do not significantly alter ring formation. Rings form when the initial bundling is effective with a longer residence time for VASP to crosslink filaments (specific binding and unbinding rates), otherwise shells form. Our predictions of effective actin bundling as a determinant of shells versus rings were validated using a mutant of VASP that has decreased bundling capability and effectively lower residence time. Finally, the shape transition of VASP droplets from spheres to ellipsoids to rods, driven by ring formation, is determined by the maximum filament length. Thus, our study identifies how different kinetic parameters can determine the actin network architecture and consequently VASP droplet deformation. From a biological perspective, this work identifies specific physical mechanisms by which protein condensates could influence the architecture of the cytoskeleton.

## Results

We begin by constructing a minimal computational model to study how the condensates affect actin organization within droplets. The model incorporates the dynamics of actin filament elongation and VASP-driven bundling. Specifically, we sought to identify the conditions in which actin can form shells and rings within VASP droplets, as observed in experiments (Figure 1A), (5). We assume that the VASP droplets have already formed and are characterized by a crowded environment with a high-density of VASP tetramers. Experiments showed that polymerized actin remains confined inside the droplet (5). Therefore, we simulated actin networks inside droplets with a high surface tension limit using non-deformable droplet-mimics characterized by a hard wall boundary condition. We used CytoSim (55, 57), an agent-based simulation platform designed to simulate cytoskeletal networks. Actin filaments are represented as a series of cylinders, each of length L_seg_ and with a flexural rigidity determined by the persistence length of actin filaments (58). Actin filaments are allowed to elongate at a predetermined rate (Figure 1C). Each of the four binding sites on the VASP tetramer can stochastically bind actin filaments that are within a binding distance of 30 nm, depending on the specified mesoscopic binding rate, k_bind_ (Figure 1D). Actin-VASP unbinding is modeled as a force-sensitive rate, given by k_unbind_. Given the stochastic nature of the simulations, we conducted multiple replicates per condition. Using this simulation platform, we systematically investigate how different biochemical and mechanical parameters can impact the architecture of the actin network in VASP droplets.

**Figure 1.**
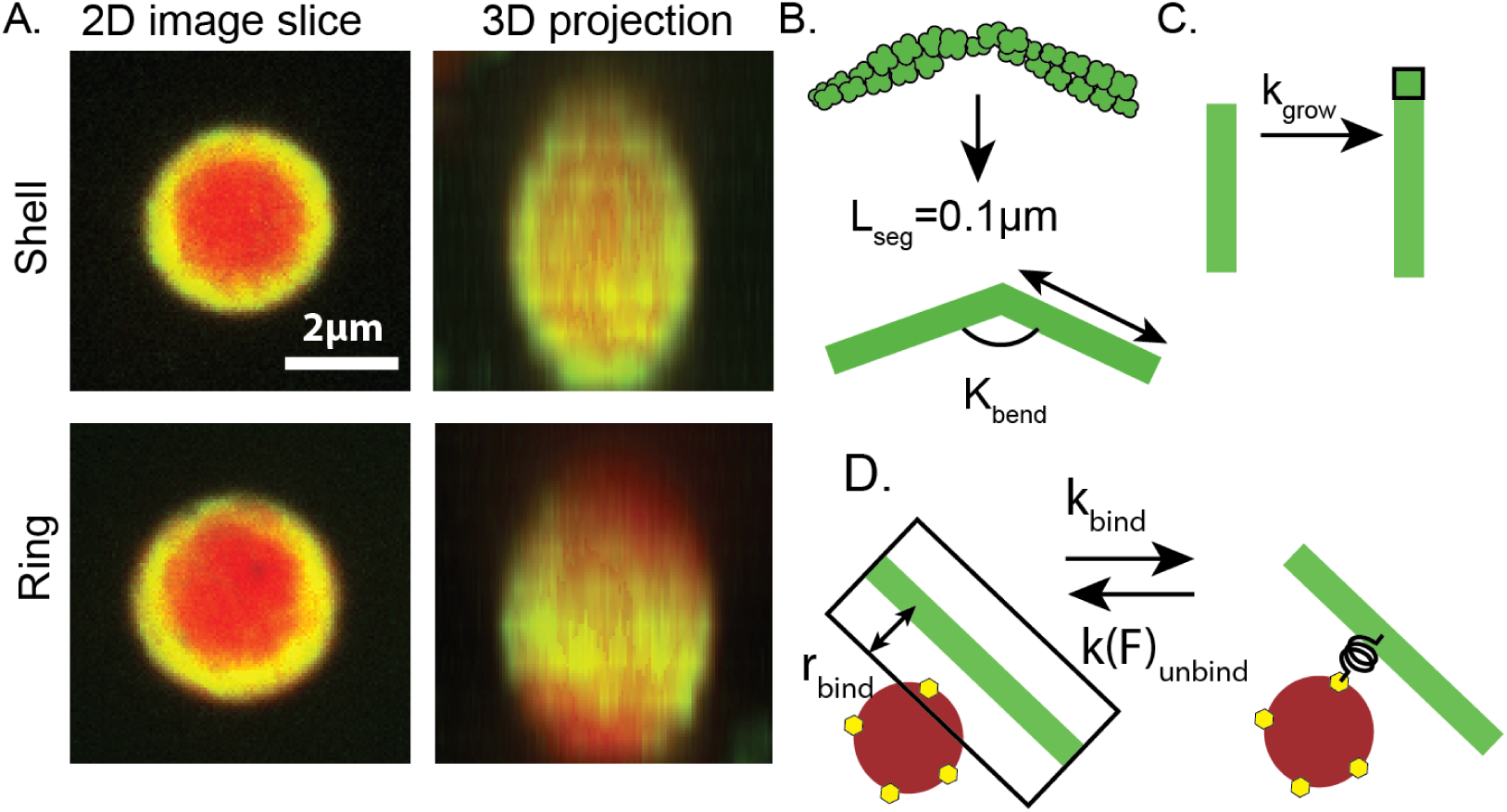
Computational model to study actin organization in VASP droplets. (A) 2D confocal image slice, and corresponding and 3D reconstructions from experimental images showing a VASP droplet with a shell- and ring-shaped actin networks from experiments. VASP is in red and actin is in green. Images show the merge of both channels. Scale Bar 2 μm. (B)-(D) Simulation set up in Cytosim for actin filaments. The VASP droplet is represented as a spherical reaction volume with hard wall boundary conditions to mimic the high surface tension limit. (B) The actin double helix is represented as a series of cylinders, each of length L_seg_ and with a flexural rigidity determined by the persistence length of actin filaments (58). (C) Filaments are allowed to grow deterministically with extension rate k_grow_. (D) VASP tetramers are represented as spherical crosslinkers with four binding sites. When a binding site encounters a filament segment within the binding distance r_bind_, a stochastic binding reaction occurs with probability determined by the mesoscopic binding rate k_bind_. Additionally, a force-sensitive unbinding reaction is also included to be consistent with detailed balance. All parameters are given in the supplementary material.

### The binding-unbinding kinetics between actin and VASP determines actin network organization in VASP droplets

We begin by simulating a spherical volume (R_drop_=1 μm) with a tetrameric crosslinker concentration of 0.40 μM (volume fraction of tetrameric crosslinker in the droplet is 2.70%). In this volume, 30 seed actin filaments, each of length 0.1 μm are distributed at random orientations throughout. The filament elongation rate is set to 10.3 nm/s such that at 600 s, the final length of actin filaments is the cross-section circumference of the droplet, 2π μm (final actin concentration is 27.67 μM and volume fraction is 3.2 x 10^-2^ %). The simulation time of 600 s was chosen based on experimental observations (5). With these initial conditions, we varied the binding and unbinding rate of VASP to actin. The dissociation constant of VASP-actin was measured to be 1.8 nM (14). However, how this dissociation constant alters the dynamic evolution of the actin network in VASP droplets, particularly due to the multivalent nature of VASP-tetramers binding to actin filaments, remains unclear. Therefore, we varied k_bind_ and k_unbind_ from 1 x 10^-4^ 1/s to 10 1/s in our simulations (Figure 2) for a fixed concentration of actin and tetramers.

**Figure 2.**
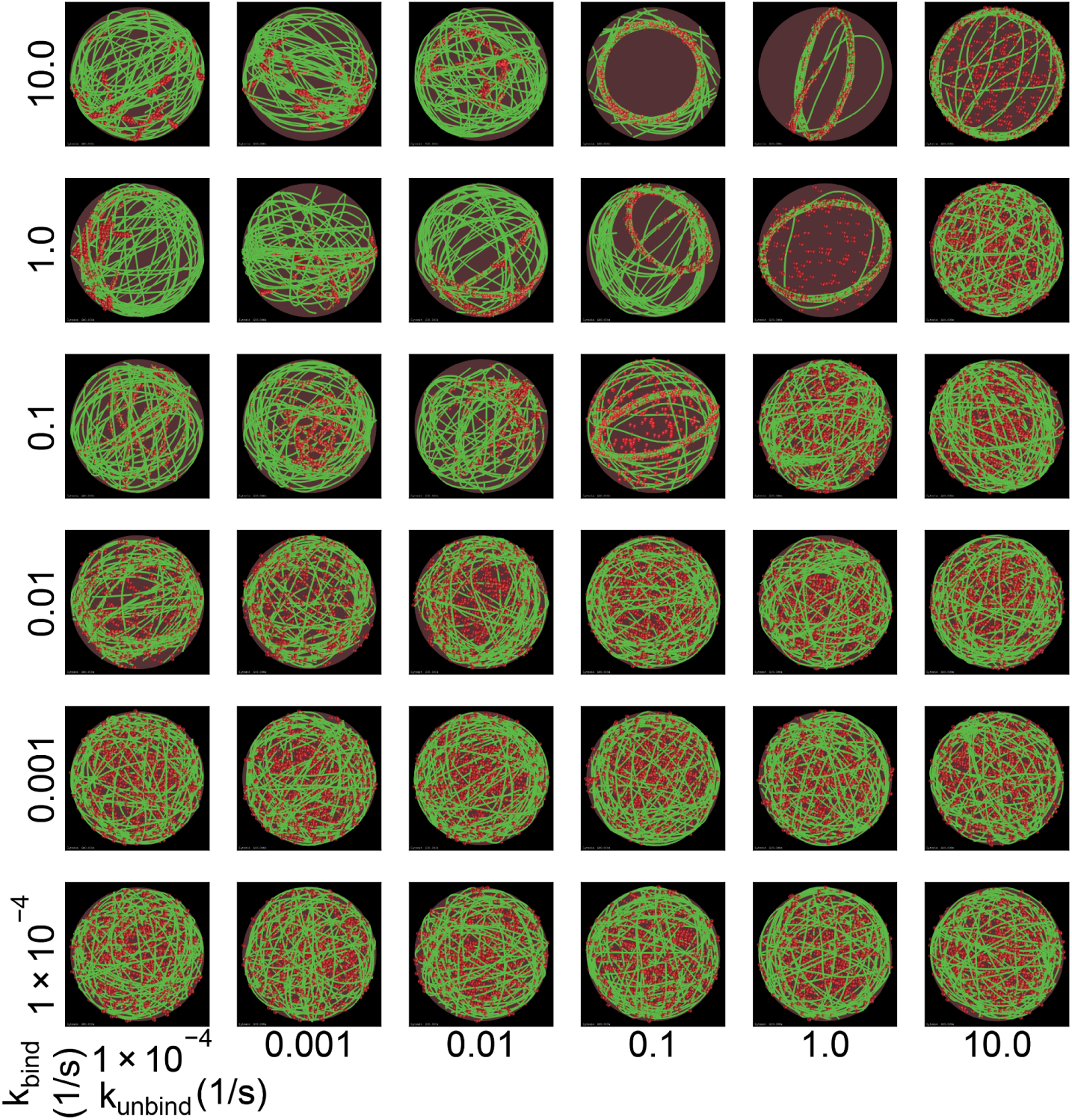
Varying crosslinker kinetics generates experimentally observed actin network shapes. Representative snapshots at t = 600 s from simulations with 30 seed actin filaments are shown corresponding to different crosslinker binding rates (k_bind_, varied along each column) and unbinding (k_unbind_, varied along each row). Actin filaments are shown in green while the tetrameric crosslinker is shown as red spheres; also see Supplementary Movie M1.

Our simulations reveal that when k_bind_ (Figure 2 rows) and k_unbind_ (Figure 2 columns) are varied, the resulting actin networks resemble experimentally observed networks such as shells, rings in addition to networks that share features of rings and shells (Figure 2). This result predicts that the crosslinking timescale determines the network organization of confined actin networks. For the combination of parameters investigated, the formation of shell like structures is more likely. The formation of VASP-decorated actin rings is favored when k_bind_ is high (1 and 10 1/s) and k_unbind_ is intermediate (1/s). Specifically, if k_unbind_ is too high, then actin filaments will not be sufficiently bundled to form a ring. In contrast, if k_unbind_ is too low, then the initial, unaligned contacts will be unable to rearrange into an aligned bundle configuration. Thus the actin shape accessibility is controlled by the time scale of actin-VASP interaction. Additionally, we note that the plane in which the ring is oriented is random in the simulations but determines the plane in which the droplet deforms (see Figure 7) (5).

Next, to quantify the actin architecture under these different conditions, we calculated the fraction of spherical surface occupied by actin (Figure 3A-B, Supplementary Figure S1). At the final timepoint, we expect actin rings to occupy a lower surface area fraction of the boundary and shells to occupy a higher surface area fraction of the boundary (Figure 3A). Calculating the kinetics of area fraction over time, we found that for all combinations of parameters tested, the initial kinetics are fairly similar for all conditions for up to 200 seconds. This is understood by recognizing that at t>184.47s, the filament reaches length >2R_drop_ and then starts to bend (Supplementary Figure S2). At low binding rates (≤0.001/s), for all values of unbinding rates tested, we found that the surface area fraction continued to grow with time, consistent with the shell-like actin organization in Figure 2. At higher binding rates, we found that for certain values of unbinding rates (for example, k_bind_=1/s, k_unbind_=10/s), the actin network formed shells at later time, while other area fraction trajectories seemed to stabilize to a smaller area fraction value (for example, k_bind_ =1/s. k_unbind_ = 1/s). These trajectories can be mapped to ring-like actin networks. We also observed that a few trajectories appeared to settle between shells and rings, suggesting that there are other intermediate structures that appear in the actin network. The strong dependency of these trajectories on kinetic rates suggests that the architectures of actin networks within VASP droplets represent kinetically trapped states.

**Figure 3.**
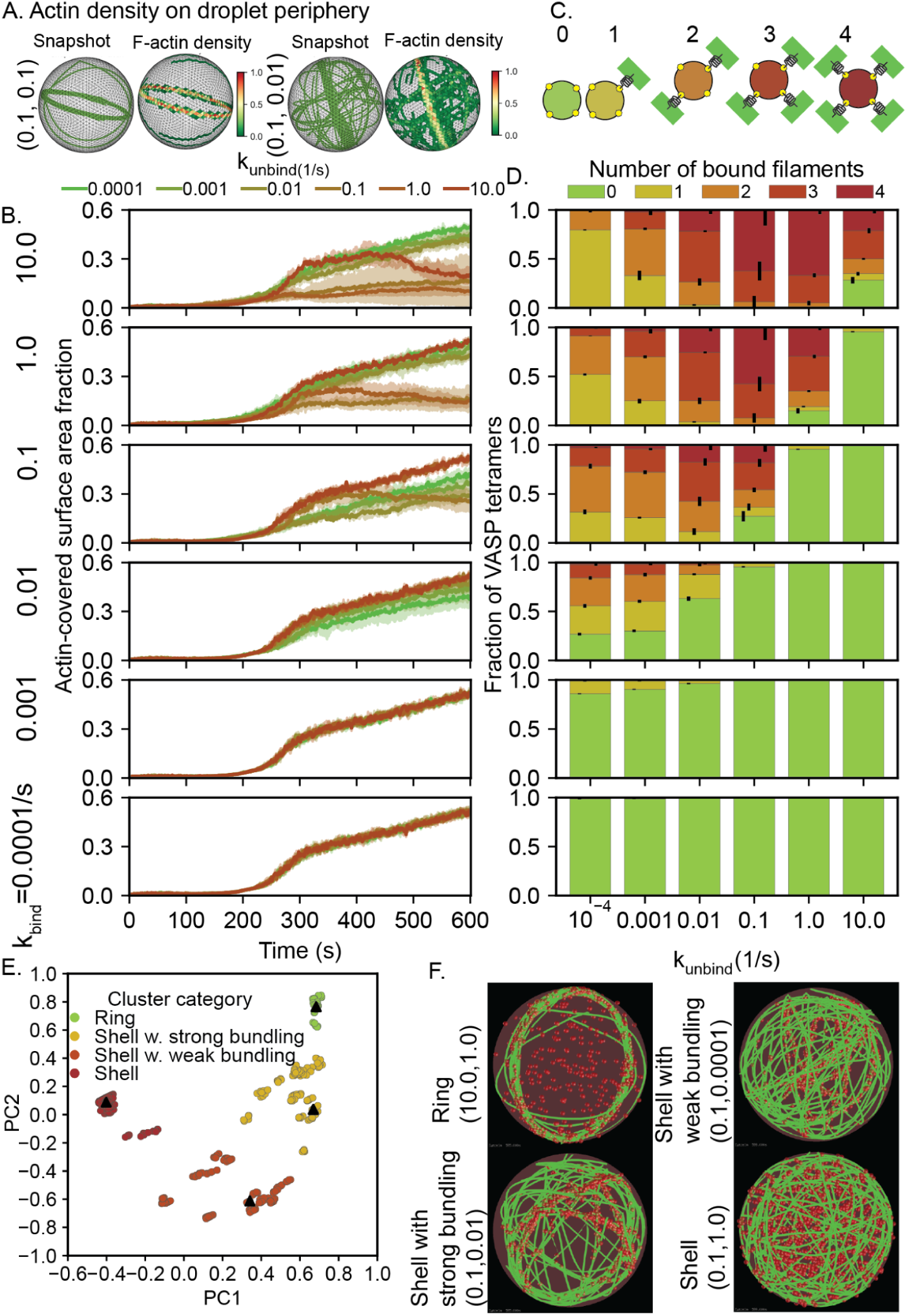
Signatures of different actin network configurations. (A) Representative final snapshots corresponding to two different kinetic parameters (given as (k_bind_, k_unbind_) to the left) are shown along with the surface actin density along the icosphere. The triangulated surface is colored based on local actin abundance (normalized). (B) Each panel shows the kinetics of the surface area fraction of the sphere covered in F-actin. The binding rate is changed across panels while each panel shows the role of VASP unbinding rate.(3 replicates per condition) (C) Cartoon illustrating the multivalent nature of VASP tetramers. (D) Each panel shows a stacked bar graph illustrating the fraction of VASP tetramers bound to 0, 1, 2, 3, and 4 filaments for different VASP unbinding rates. Corresponding VASP binding rate is shown to the left of panel B and the errorbar shows the standard deviation. For (B) and (D) data was obtained from the last 5% of each of the five replicate trajectories (30 snapshots). (E) 5 snapshots per replicates were considered from each of the 35 kinetic parameters studied in Figure 1 and the data shown in panels B and D were used to cluster data using K-means clustering. The resulting 4 clusters using the first 2 principal components (PCs) are shown in (F). along with the snapshot (identified by the k_bind_, k_unbind_ pair) closest to each cluster centroid (solid triangles).

To further classify these networks, we calculated the extent of filament bundling (see Supplementary Material for details). The extent of bundling was quantified by counting the number of crosslinkers that are bound to 0 (unbound), 1, 2, 3, or 4 filaments (Figure 3C-D). We found that conditions that formed shell networks are associated with poor bundling and have a high number of unbound crosslinkers (Figure 3D, left column) because the residence time of crosslinkers is very low. On the other hand, ring networks have a higher fraction of crosslinkers bound to at least 3 filaments (Figure 3D right column). Finally, shell-ring intermediates have a significant number of crosslinkers bound to more than 2 filaments.

Thus far, our analysis has been based on visual inspection of the actin networks and corresponding metrics (area fraction and non-zero number of filaments bound to a crosslinker, a total of 5 order parameters). These five dimensions are not independent but correlated to some extent (Supplementary Figure S3). We reduced the dimensionality of the system using principal component analysis and found that the first three principal components (PCs) (59) explained 97.64% of the variance of the entire dataset (Figure S3).To interpret the relative contributions of the original five dimensions on each of the PCs, we employed varimax rotation to calculate the loadings. We found that each of our five order parameters contribute strongly to at least one of the three PCs. Additionally, we observe from the varimax loading (Figure S3) that PC1 is positively correlated with the number of crosslinkers bound to 3 and 4 filaments. Additionally, it is negatively correlated with surface area fraction. Plotting the two PCs, we see that shells have a low coordinate along PC1 consistent with both a high surface area-fraction, and the low number of VASP molecules bound to 3 and 4 filaments. Therefore, we hypothesized that the various actin shapes in our dataset should be localized to distinct regions of the PC space. Employing K-means clustering analysis along the first three PCs, we identify four distinct clusters in our dataset. (Figure 3E, F, S3). Visual inspection of snapshots within each cluster reveals that shells and rings form two distinct clusters while the other two clusters consist of shells with weak bundling and shells with strong bundling (Figure 3E, F). To understand the shape differences between the clusters, we computed the moment of inertia matrix for each of the actin shapes and calculated the spans approximating the network shape as an ellipsoid. The distribution of ellipsoidal spans are plotted for each of the clusters in Figure S3 and we see distinct differences in shapes between each of the four clusters identified. Grouping the snapshots based on their k_bind_, k_unbind_ values, we find that in a subset of conditions, the emergent actin shape is probabilistic (Supplementary Table S2). These findings predict that a nontrivial combination of actin-VASP binding-unbinding rates and number of filaments bound to each VASP molecule determine the network architecture.

### Actin ring formation is robust to changes in filament elongation rate

We next asked if the kinetics of actin elongation could affect the probability of ring formation. We conducted two separate sets of simulations to answer this question – we varied the final filament length (Figure 4) and the actin elongation rate (Figure 5). These two variations were designed based on the knowledge that VASP promotes actin filament elongation as a processive polymerase and the polymerization rates can depend on the actin organization itself. VASP is known to gather and promote elongation of multiple barbed ends simultaneously and VASP-mediated polymerization rates of shared barbed ends can be up to 3 times faster than free barbed ends (14).

**Figure 4.**
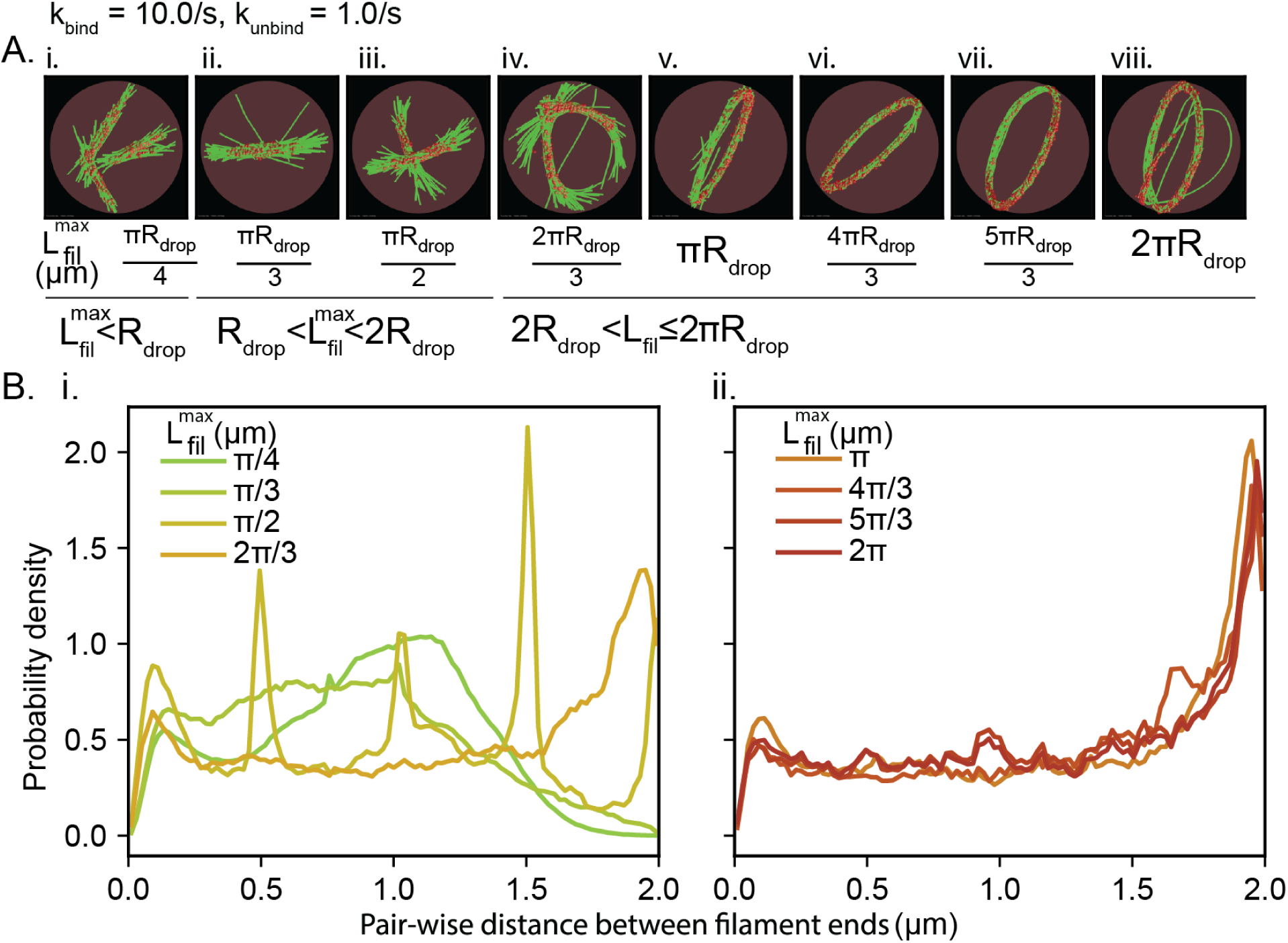
Actin networks with growing filaments form rings when final filament length is greater than the diameter of the droplet. A. Representative final snapshots (t=600s) from simulations under ring-forming conditions at various final filament lengths; also see Supplementary Movie M2. B. Probability density functions of pairwise distances between filament ends is shown under various L_fil_^max^. (Data used: Final 5 snapshots from each of the 5 replicates per condition)

**Figure 5.**
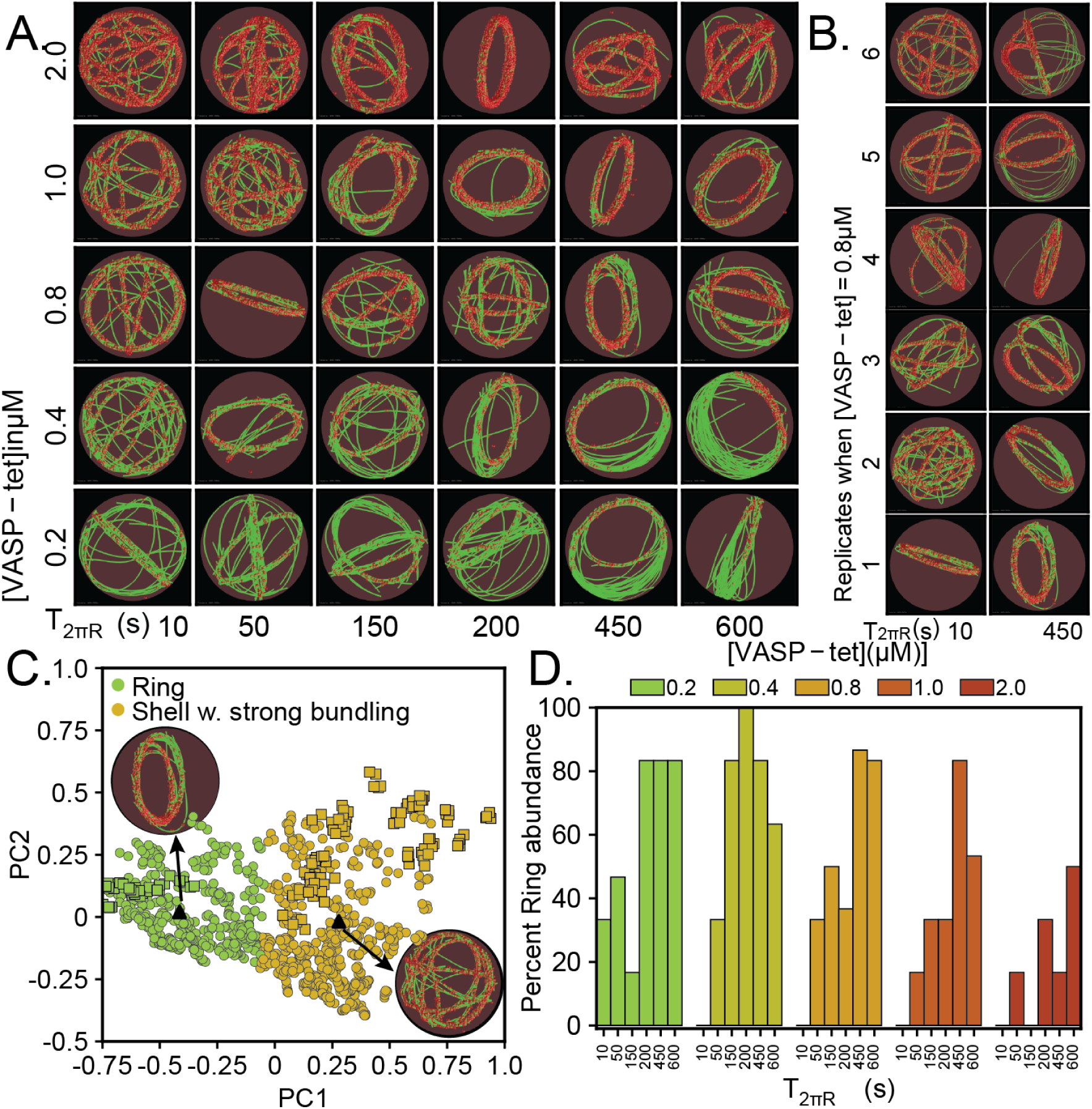
Simulations predict slower elongation rate (high T_2πR_) increases ring formation probability. A. Representative final snapshots (t=600 s) from simulations at various [VASP-tet] and elongation rates measured by T_2πR_, the time for filaments to grow to 2πR_drop_ length (k_bind_=1.0/s, k_unbind_=0.1/s); also see Supplementary Movie M3. Rings are more probable when T_2πR_ is large. B. We find that the final network shape has higher heterogeneity at low T_2πR_ (data shown for tetramer concentration of 0.8 μM from 5 replicates). C. Simulation results from this study (circle) were combined with data from Figure 3E (shown as squares) and clustered to identify salient network shapes. Combined data projected along the first two principal components are shown, colored by cluster. The datapoint from the current study that are closest to each of the cluster-centroids are shown as triangles along with insets of the final snapshot. Supplementary Figure S7 shows details of clustering algorithm and Figure S8 shows the distribution of ellipsoidal span for rings and rings with strong bundling D. Probability of ring formation calculated from the clustered data is shown as a function of [VASP] and filament elongation timescale. Data used: Five snapshots from each of the 10 replicates. Please refer to Supplementary Figure S4 for plots of actin network shapes in each of the four categories and Figure S6 for

We fix the k_bind_, k_unbind_ values where we know rings will form (10s^-1^ and 1s^-1^) and change the maximum filament length, L^max^_fil_ that can be attained in the simulations at 600 s (Figure 4). We explore the role of droplet diameter and circumference in controlling actin shape by simulating L^max^_fil_ <R_drop_, R_drop_<L^max^_fil_ <2R_drop_, 2R_drop_<L^max^_fil_ <πR_drop_, and πR_drop_<L^max^_fil_ <2πR_drop_. The number of filaments and actin addition rate were calculated so that the rate of F-actin addition remains constant. Thus, in this simulation, we are effectively changing the elongation rate of individual filaments by controlling the final filament length. For all filament lengths, we see that actin forms bundles, consistent with the role played by the binding and unbinding rates. When the maximum filament length is smaller than the radius of the droplet, we see that the bundles are linear (Figure 4A i-ii) and probability distribution of the end of end distances of filaments peaks at distances smaller than the droplet diameter (Figure 4B i). When the filament length is greater than the radius of the droplet, but less than the diameter of the droplet, we observe some bending of the rings (Figure 4A iii-iv) and the peak distribution shifts closer to the diameter but not quite towards rings (Figure 4B i). When the filament length is greater than the diameter of the droplet, we observe that the filament bundles always form rings (Figure 4A, v-viii) and the end-to-end distance probability is close to the droplet diameter (Figure 4B ii).

Next, we fixed the final filament length and varied the filament elongation rate. We define the time it takes a filament to grow to length 2πR_drop_ (one circumference along the droplet) as T_2πR_. We vary T_2πR_ between 10 s and 600 s (Figure 5). To understand the role of T_2πR_, we chose binding parameters where ring formation was probabilistic (k_bind_ =0.1/s, k_unbind_=0.1/s). We also varied the VASP concentration to investigate if it had a role to play in ring formation. We observed that actin rings can be obtained for a number of combinations of elongation rate and VASP concentration (Figure 5A). Furthermore, we observed that for a fixed VASP concentration (Figure 5B, showing VASP = 0.8 μM, see Supplementary Figure S6 for others), the network configuration was more heterogeneous across replicates at lower values of T_2πR_ (faster elongation) than at higher values of T_2πR_ (slower elongation). To quantify the variation in the actin networks, we plotted the principal components by combining the earlier dataset (Figure 3) with the current dataset. We found that despite the apparent differences in the network architecture, only two possible network configurations were seen – actin rings and shells with strong bundling (Supplementary Figure S5). We repeated the clustering and principal component analyses using an updated dataset with data points corresponding to the ring and shells with strong bundling from the earlier dataset and all data points of the current dataset (Figure 5C). Finally, we found that while the ring probability increases with slower elongation rate (larger T_2πR_), the remainder of the probability is to form shells with strong bundling (Figure 5D). Thus, we find that actin ring formation probability is robust to changes in actin elongation rate as long as the k_bind_ and k_unbind_ are favorable for ring formation, suggesting that actin-VASP interaction timescale dominates ring formation.

### Weakening filament bundling results in loss of rings

Thus far, our simulations predict that ring formation occurs when the initial bundling is effective (large effective residence time for actin-VASP multivalent binding), and that ring formation is robust to changes in actin elongation rates. If this is true, then decreasing the residence time of VASP tetramers on actin filaments should change the kinetic trajectories from rings to other network configurations. To test this hypothesis, we choose a k_bind_, k_unbind_ pair where rings form (1, 1) and slowly increase k_unbind_ (Figure 6A). Increasing k_unbind_ decreases the effective residence time of VASP on actin filaments and therefore its effective bundling capability. Indeed we observed that the configuration of the actin network changed from rings to shell-ring intermediates (Figure 6A). Further quantification of the distribution of actin network architectures as k_unbind_ increases shows that weakening bundling by increasing k_unbind_ shifts the abundance of actin networks from rings and shells with strong bundling to shells (Figure 6B).

**Figure 6.**
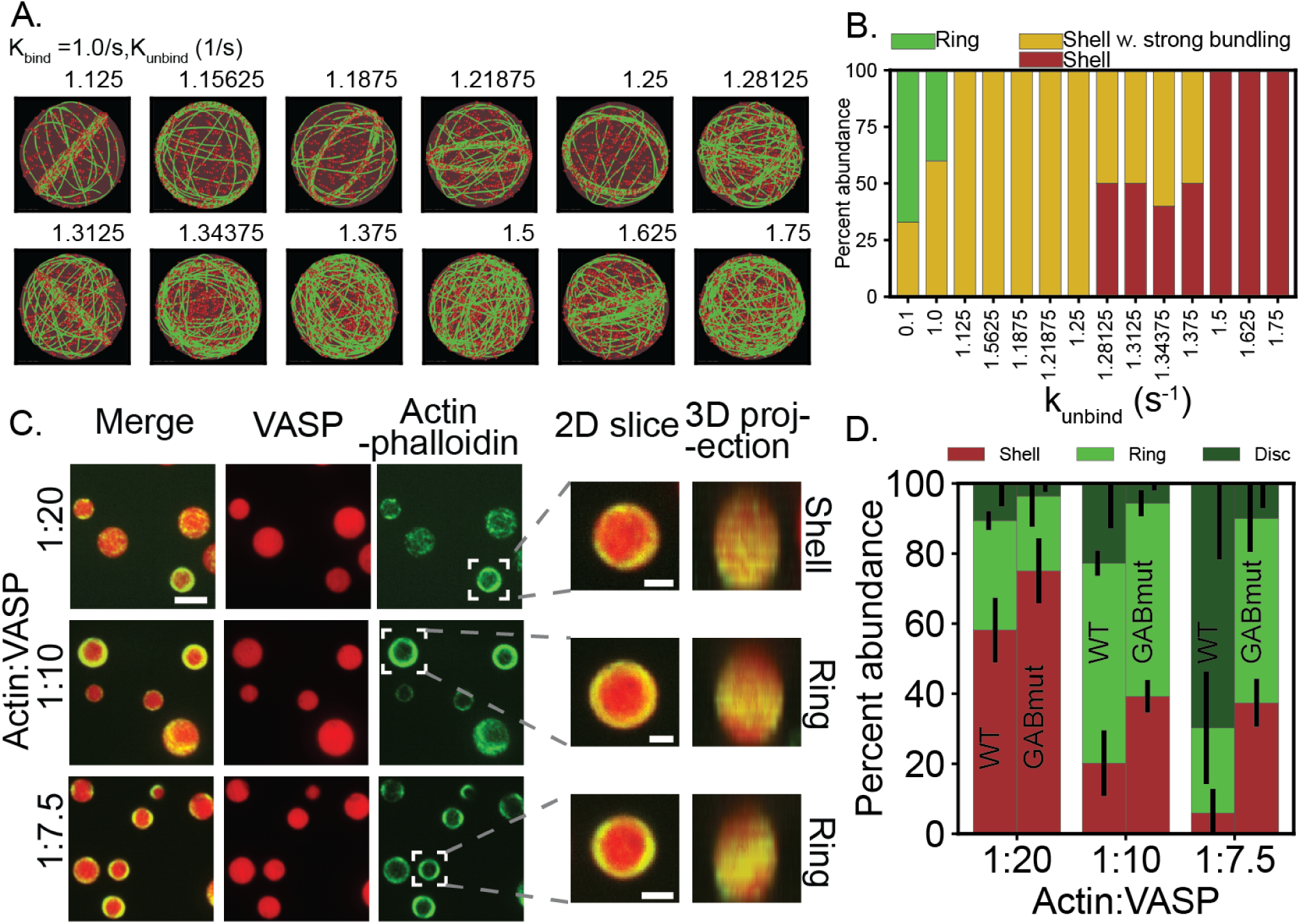
VASP-actin dissociation constant plays a key role in determining the actin network shape. (A) Representative final snapshots (t=600) obtained from simulations for different k_unbind_ values for k_bind_ =1.0/s. (B) The resulting final actin shapes (N = 10 for Kd = {1.28125,1.3125,1.34375}, N = 5 otherwise), were used to compute probability of observing ring-like and shell-like actin. Please refer to Supplementary Figure S9 for details of PCA and clustering. (C) Experimental images showing VASP-GABmut droplets containing phalloidin-stained actin at various Actin:VASP ratios. Insets show selected droplets with peripheral actin accumulation that were classified as shells or rings from 3D projections. Overview scale bar = 3 μm; Inset scale bar = 1 μm. (D) Experimental data shows mean and standard deviation of percentage abundance corresponding to Shell, Ring, and Discs for WT and GABmut-VASP (N= 3 biologically independent experiments).

In order to test our prediction that a combination of actin-VASP dissociation kinetics determines the network architecture, we sought to mutate VASP in ways that reduce its association to actin filaments. The most obvious way to disrupt actin-VASP interaction would be to mutate VASP’s F-actin binding site. However, this mutation is too severe to be useful in the current study, as it inhibits bundling of filaments so strongly that peripheral accumulation of actin filaments in VASP droplets is completely abolished (5). In contrast, mutation of VASP’s G-actin binding (GAB) site has a more modest impact on filament binding and bundling (34, 36), such that it provides a useful tool for studying the effects of impaired bundling on the abundance of shells, rings, and discs. To test this idea, we formed droplets of VASP-GABmut, as previously described (5), and added unlabeled actin monomers to them. We then stained with phalloidin to specifically visualize actin filaments and performed confocal fluorescence microscopy to determine the spatial arrangement of the filaments within the droplets. Upon three dimensional reconstruction of the confocal slices, droplets that contained actin shells and rings, along with droplets that deformed into discs, were observed (Figure 4C). We quantified the abundance of the shells, rings, and discs as a function of actin to VASP ratio (Figure 6D). Using droplets composed of wild-type VASP, we have previously shown that actin filaments adopt increasingly bundled structures as actin to VASP ratio increases. This increase in bundling results in a higher abundance of rings and discs, and a corresponding decrease in shells (data from (5) replotted). However, using VASP-GABmut, we found that, as the actin to VASP ratio increased, there were substantially fewer droplets that contained bundled actin structures. Specifically, at an actin to VASP ratio of 1:7.5, a majority of droplets composed of wild-type VASP had deformed into discs (70 ±13% mean±SEM), while less than 10% of droplets composed of VASP-GABmut deformed into discs, with the majority consisting of spherical droplets that contained actin rings (53±6% mean±SEM) (Figure 6D). Similarly, as the actin to VASP ratio increased for droplets composed of wild-type VASP, fewer contained actin shells, with the majority containing actin rings. In contrast, more than a third of droplets composed of VASP-GABmut contained actin shells, even at an actin to VASP ratios of 1:7.5 (37±4% mean±SEM). Overall, these data suggest that the consequence of reduced filament bundling by VASP-GABmut is a reduction in the number of bundled actin structures, specifically rings and discs, compared to droplets composed of wild-type VASP. Collectively, simulations and experiments confirm that the formation of actin rings requires strong bundling and large residence time of VASP on actin.

### Actin bundle reorganization determines the maximum aspect ratio of the VASP droplet

Previously, we showed that when actin rings reach a critical thickness, droplets deform and become ellipsoidal. We also showed that at high actin to VASP ratios, the droplet assumes a rod shape, which contains a linear actin bundle (Figure 5 in (5)). Therefore, we next investigated the features of actin rings that determine the aspect ratio of VASP droplets. We begin these simulations with a spherical droplet in which a ring has already formed. To mimic the deformation of the droplet, we change the sphere to a spheroid by changing the major axis and minor axis 10 nm at a time. We then let the system relax into the new droplet geometry for 1s. In this process, we assume that the relaxation of actin shape results from a combination of mechanical dissipation of excess bending energy and mechanochemical reorganization of crosslinkers. We observe that for a fixed value of maximum filament length equalling πR_drop_, as we deform the droplet, the actin ring continues to track the surface of the droplet, and at high aspect ratios, the ring unravels to become a linear bundle (Figure 7A). Thus, we mimic the sphere-to-ellipsoid and ellipsoid-to-rod transitions of the droplet shapes. We next varied the maximum filament length and calculated the allowed maximum aspect ratio of the droplet (Figure 7B). The maximum aspect ratio is defined as the droplet shape at which the droplet ends still have adequate actin to remain in contact (Supplementary Figure S11). The maximum aspect ratio is shown by the solid black line when the major axis of the ellipse equals the maximum final filament length (Figure 7B, C). We observe that at low maximum filament lengths, the droplets deform to aspect ratios higher than the maximum aspect ratio. This is because the filament bundles are pliant, and filaments are able to slide past one another to create a bundle longer than the maximum filament length. However, we observe that there is a threshold value of maximum filament length beyond which the aspect ratio is lower than the maximum permitted value. This is because at higher L_fil_ ^max^ values, bundles no longer reorganize and the rings remain kinetically trapped. Thus, we find that while the sphere-ellipse transition occurs under a wider range of ring-forming conditions, the maximum filament length plays a critical role in determining where the ellipse to rod transition will take place.

**Figure 7.**
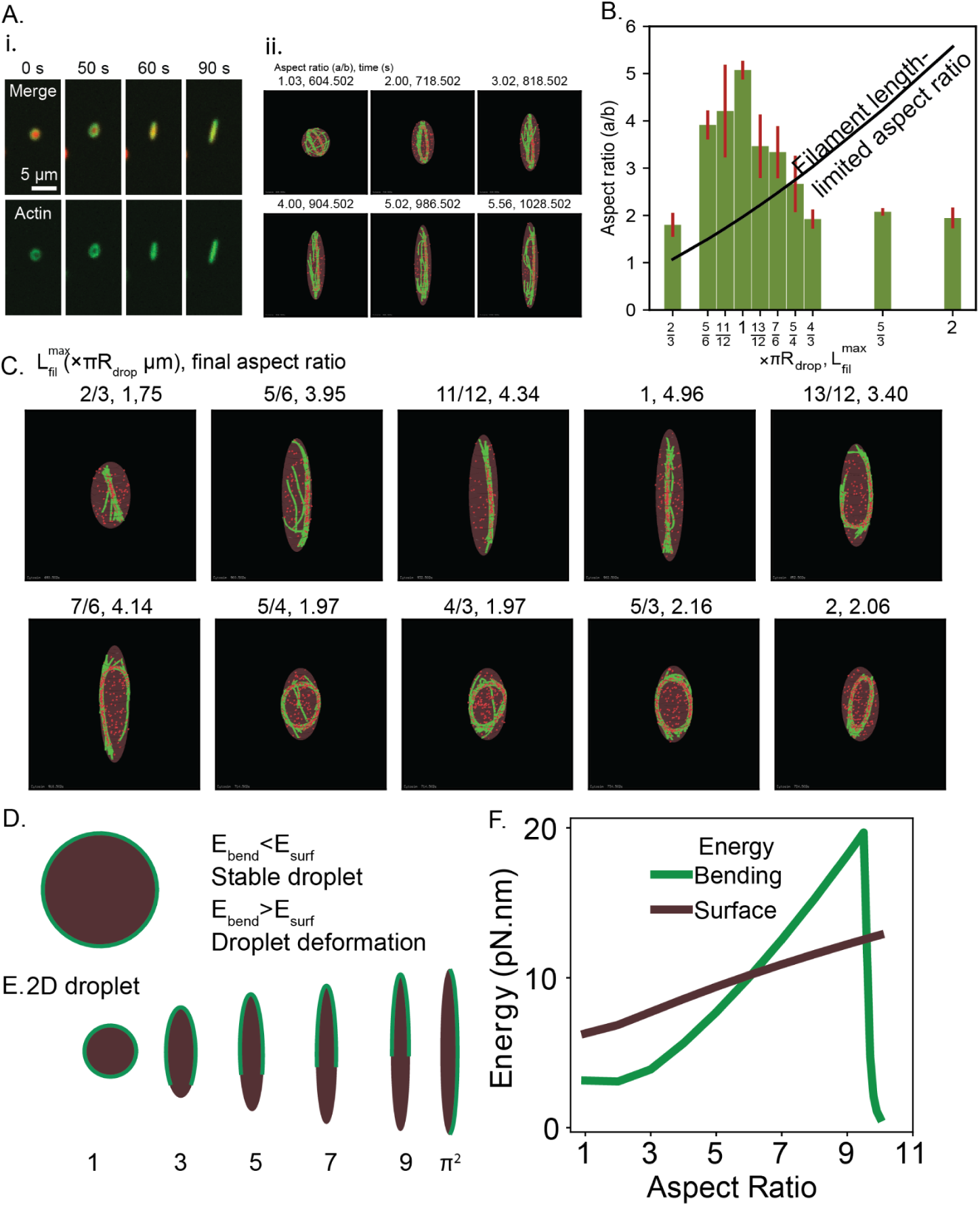
Filament length determines the maximum aspect ratio of the droplet. A. i. Linear droplets that elongate over time are formed by addition of 2 μM actin to 10 μM VASP droplets Scale bar 5 μm ii.Time series of snapshots corresponding to L_fil_^max^ = πR_drop_ shows the transition of a network with rings to an ellipsoidal droplet with rings and eventually, extended bundle. Please see Supplementary Movie M4 for a video of trajectories. B. Bar plot of mean and standard deviation in the maximum aspect ratio of droplets at various L_fil_^max^ values (5 replicates). The maximum aspect ratio was determined by the protocol outlined in Supplementary Methods to ensure filaments remain in contact with the high-curvature tips of the ellipsoid. Solid line shows the maximum aspect ratio permitted when the ellipsoidal span, 2a = L_fil_^max^. C. Representative maximum aspect ratio snapshots from simulations with different L_fil_^max^. Actin filaments (green) and VASP tetramers(red spheres) are shown within droplets (brown). D. Energetic considerations causing droplet deformation. E. Droplet deformation is driven by the competition between actin bending energy and droplet surface energy. F. Bending and surface energy as a function of droplet aspect ratio.

## Discussion

Many actin-binding proteins are thought to undergo liquid-liquid phase separation (60). The role of these phase-separated domains in organizing actin networks has been studied in different systems. Recent studies have shown that abLIM1 condensates induced asters and webs of F-actin filaments in vitro (4). Phase separation of Nephrin/Nck/WASP on lipid bilayers increases the dwell time of N-WASP and Arp2/3 complex, increasing actin assembly (3). Previously, we showed that VASP phase separates in liquid-like droplets and that within these droplets, actin can adopt a number of configurations including shells and rings. We also showed that VASP droplets will deform when actin rings exceed a certain thickness (5).

In this study, we used computational modeling to simulate the interactions between actin and VASP to mimic experimental observations of actin polymerization in a phase-separated VASP droplet. Our simulations revealed that the actin network can become shells, shells with weak bundling, shells with strong bundling, and rings. All these structures form due to stochastic ratcheting of VASP where VASP molecules stabilize the current actin configuration and also steer the network shape changes in the next instance. As a result, binding of several VASP molecules imposes energetic barriers on drastic reorganization of actin shape due to kinetic trapping. This ability to overcome barriers is less probable in the experimental timescales. We then focused on the formation of actin rings, which are responsible for droplet deformation. We found that rings that promote deformation of the spherical droplet are made of filaments that are below a certain critical length; above that length, the rings remain trapped in the bundled state and do not straighten out to further deform the droplet. These results may explain the experimental observation that for droplets with the same actin concentration, some deform into rods but some remain stuck with rings in them (See Figure 5b of (5))

The sphere-to-ellipsoid transition and the ellipsoid-to-rod transitions are consistent with our previous energy balance model presented in (5). The first transition from a sphere to a spheroid is consistent with our previous 2D model for droplet deformation, where, circular droplets deform into elliptical droplets when the actin ring bending energy is greater than the droplet surface energy (Figure 7D). We gain additional novel insights from the explicit consideration of bundle reorganization. We show that actin networks can form rings over a range of filaments lengths. In addition, consistent with experiments, we find that the ellipsoidal droplets can take a wide range of aspect ratios because of the dependence on bundle thickness. For the ellipsoid to rod transition, we show that the bundle needs to reorganize by unfolding its curvature, which is consistent with the two energy barriers we identified previously (Figure 7E, F). Additionally, we see that filament length plays a critical role in aiding the unfolding. Curvature unfolding requires extensive reorganization of the bound VASP crosslinkers to allow for bending energy relaxation of actin filaments. Rings with filaments L_fil_ ^max^∼πR deform readily by unfolding resulting in a bundle longer than πR_drop_ and therefore surpassing the filament length-limited aspect ratio. When the filaments are shorter than the ring circumference, the stabilization of ring shape is aided by actin-VASP contacts. Upon droplet shape change, VASP molecules undergo force-sensitive unbinding to dissipate excess mechanochemical energy resulting in ellipsoid-rod transition. On the other hand, rings made of filaments L ^max^>>πR cannot reorganize themselves to reach the filament length-limited aspect ratio. Because the filaments forming such rings trace a significant portion of the ring circumference, the resulting rings resist deformation due to steric-driven restriction on filament unfolding and constant reorganization of actin-VASP contacts. When deformed, such bundles form stable, elliptical rings. Thus, we demonstrate that the internal degrees of freedom of a bundle play a critical role in droplet shape change.

The stabilization of rings with L ^max^>>πR in the frustrated elliptical shape away from the equilibrium linear bundle can be explained by kinetic trapping. Kinetic trapping stabilizes long-lived steady-states that are far-away from the minimum energy configurations (61, 62). Kinetic trapping plays a significant role in self-assembly processes involving multi-body interactions (63–66). In our study, we see that the rings with L ^max^>>πR form elliptical rings rather than unfolding to a linear bundle configuration. The stochastic unbinding of each of the four binding sites on VASP tetramers along with the steric repulsion between actin-actin and actin-VASP molecules stabilize elliptical configurations. Thus, our results suggest that the relative abundance of rod-shaped droplets should increase at longer time scales. Importantly, kinetic trapping has been observed in various cytoskeletal systems (67, 68). Specifically, crosslinker proteins have been shown to enhance contractility of actin networks (40) and mitotic spindle assembly (69). Here, we establish additional mechanisms by which kinetic trapping might guide actin reorganization.

In summary, our work offers a potentially simple mechanism for formation of various actin shapes that are relevant for cellular function. Future iterations of such models should include explicit considerations of the viscoelastic nature of the droplet and explicit diffusion of components across the droplet interface. Incorporating such complex physical behaviors would lead to a more robust understanding of the role of such droplets in actin regulation. It would also be insightful to study formation of protein composites that include multiple actin binding proteins and their role in determining the actin network architecture. Particularly, additional nucleation in the form of Arp2/3 could help explain the filopodial initiation from lamellipodia as found in cells (70). As VASP is found close to the membrane, along with a whole host of other actin binding proteins, it is possible that interactions with other actin-binding proteins could lead to further refinement of the kinetic trapping mechanisms proposed here.

## Acknowledgments

The authors would like to thank Dr. Yossi Eliaz, Dr. Christopher Lee, Dr. Mayte Bonilla Quintana, Dr. Ashwin Ravichandran and Dr. Sriram Vignesh Mani for constructive feedback on the model and results presented here. They would also like to thank Prof. Francois Nedelec for the feedback on the model schematic. This work was supported by NSF DMS 1934411 to P.R. and J.C.S, Office of Naval Research N00014-20-1-2469 to P.R, and NIH (R35GM139531) to J.C.S.

## Supplementary Material

### Supplementary Tables

**Table S1:**
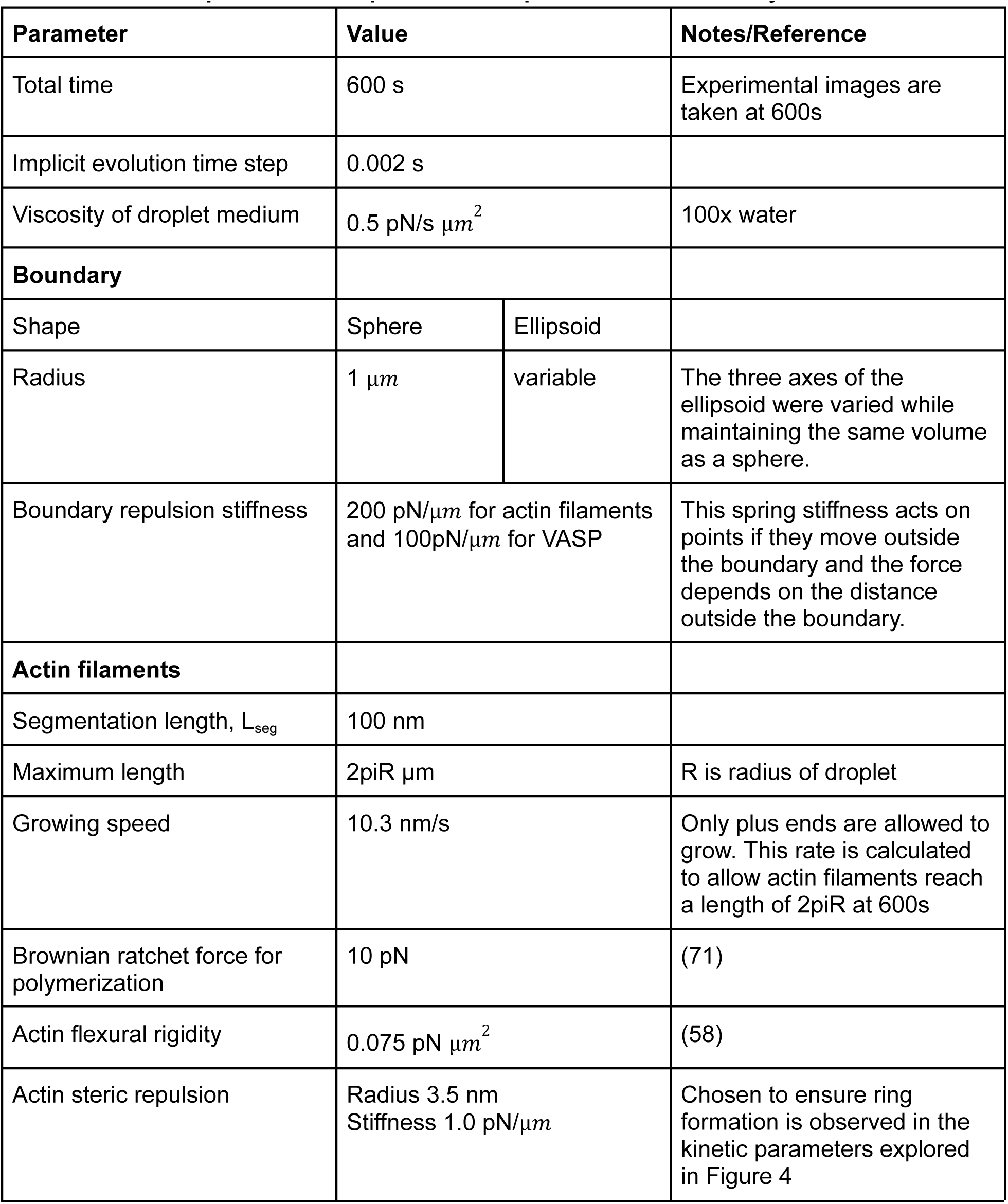
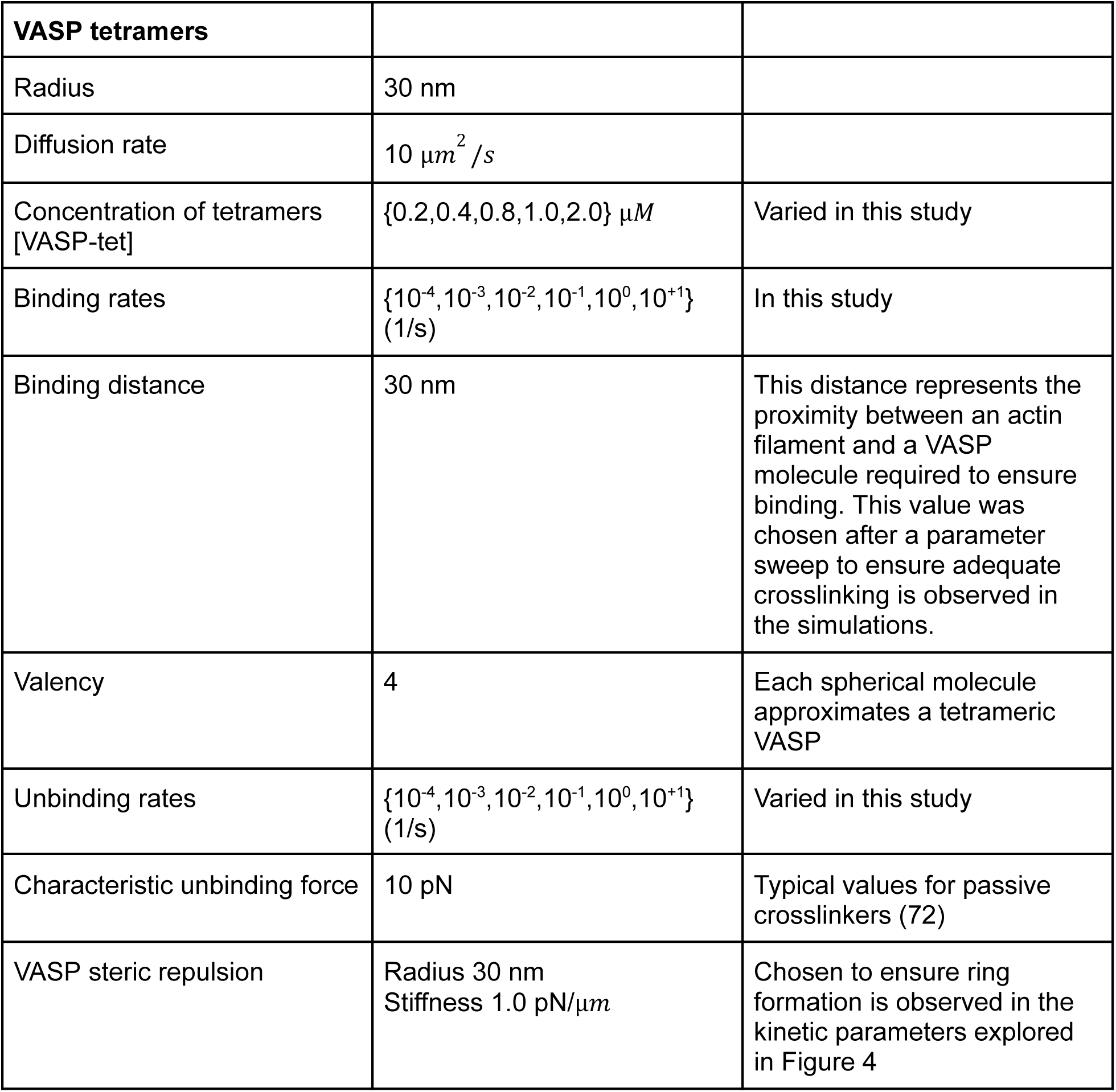
Table of parameters required to set up the actin model in Cytosim.

| Parameter | Value |  | Notes/Reference |
| --- | --- | --- | --- |
| Total time | 600 s |  | Experimental images are taken at 600s |
| Implicit evolution time step | 0.002 s |  |  |
| Viscosity of droplet medium | 0.5 pN/s $\mu m^2$ | | 100x water |
| **Boundary** |  |  |  |
| Shape | Sphere | Ellipsoid |  |
| Radius | 1 $\mu m$ | variable | The three axes of the ellipsoid were varied while maintaining the same volume as a sphere. |
| Boundary repulsion stiffness | 200 pN/$\mu m$ for actin filaments and 100pN/$\mu m$ for VASP | | This spring stiffness acts on points if they move outside the boundary and the force depends on the distance outside the boundary. |
| **Actin filaments** |  |  |  |
| Segmentation length, $L_{seg}$ | 100 nm | | |
| Maximum length | 2πR $\mu m$ | | R is radius of droplet |
| Growing speed | 10.3 nm/s | | Only plus ends are allowed to grow. This rate is calculated to allow actin filaments reach a length of 2πR at 600s |
| Brownian ratchet force for polymerization | 10 pN |  | (71) |
| Actin flexural rigidity | 0.075 pN $\mu m^2$ | | (58) |
| Actin steric repulsion | Radius 3.5 nm<br>Stiffness 1.0 pN/$\mu m$ | | Chosen to ensure ring formation is observed in the kinetic parameters explored in Figure 4 |

|  |  |  |
| --- | --- | --- |
| **VASP tetramers** |  |  |
| Radius | 30 nm |  |
| Diffusion rate | $10 \mu m^2 / s$ | |
| Concentration of tetramers [VASP-tet] | $\{0.2, 0.4, 0.8, 1.0, 2.0\} \mu M$ | Varied in this study |
| Binding rates | $\{10^{-4}, 10^{-3}, 10^{-2}, 10^{-1}, 10^0, 10^{+1}\} (1/s)$ | In this study |
| Binding distance | 30 nm | This distance represents the proximity between an actin filament and a VASP molecule required to ensure binding. This value was chosen after a parameter sweep to ensure adequate crosslinking is observed in the simulations. |
| Valency | 4 | Each spherical molecule approximates a tetrameric VASP |
| Unbinding rates | $\{10^{-4}, 10^{-3}, 10^{-2}, 10^{-1}, 10^0, 10^{+1}\} (1/s)$ | Varied in this study |
| Characteristic unbinding force | 10 pN | Typical values for passive crosslinkers (72) |
| VASP steric repulsion | Radius 30 nm<br>Stiffness $1.0 \text{ pN}/\mu m$ | Chosen to ensure ring formation is observed in the kinetic parameters explored in Figure 4 |

**Table S2.** Table of actin shapes corresponding to each K_bind_ and K_unbind_ value.

|  |  |  |  |  |  |  |  |
| --- | --- | --- | --- | --- | --- | --- | --- |
| $K_{\text{bind}} \text{ (s}^{-1}\text{)}$ | $10^{-4}$ | Shell with weak bundling | Shell with weak bundling | Shell with strong bundling | Shell with strong bundling - 33.33%<br>Ring - 66.67% | Ring | Shell with strong bundling |
| | $10^{-3}$ | Shell with weak bundling | Shell with weak bundling | Shell with strong bundling | Shell with strong bundling - 33.33%<br>Ring - 66.67% | Shell with strong bundling | Shell with strong bundling |
| | $10^{-2}$ | Shell with weak bundling | Shell with weak bundling | Shell with strong bundling | Shell with strong bundling | Shell | Shell |
| | $10^{-1}$ | Shell with weak bundling | Shell with weak bundling | Shell | Shell | Shell | Shell |
|  | 1.0 | Shell | Shell | Shell | Shell | Shell | Shell |
|  | 10 | Shell | Shell | Shell | Shell | Shell | Shell |
| $K_{\text{unbind}} \text{ (s}^{-1}\text{)}$ | | $10^{-4}$ | $10^{-3}$ | $10^{-2}$ | $10^{-1}$ | 1.0 | 10 |

### Supplementary Methods

#### Chemical and mechanical framework employed in CytoSim

Simulations were performed in Cytosim (https://gitlab.com/f-nedelec/cytosim). Cytosim is an agent-based framework to simulate the chemical dynamics of filamentous networks while also accounting for their mechanical properties (55, 57). Towards this, Cytosim numerically solves a constrained Langevin framework in a viscous medium at short time intervals (2 ms in this study) to model the dynamics of filaments along with the diffusing species. To capture essential chemical reactions observed in VASP droplets, filament extension is modeled as a deterministic force-sensitive process along with stochastic Monte Carlo sampling of crosslinker binding and its force-sensitive unbinding reactions.

#### Representation of actin filaments

Actin filaments are represented as inextensible fibers whose contour is traced by a series of linear segments each of length 100 nm connected at hinge points (Figure 1A). CytoSim computes bending energy of the fiber based on the flexural rigidity specified in the input parameters.

#### Representation of VASP tetramer

VASP tetramers are known to bind up to four actin filaments through the F-actin binding domain (35). Additionally, VASP also has a G-actin binding domain which assists in increasing the extension rates of actin filaments (73). Thus, VASP can act as an actin polymerase and a crosslinking protein (15). Given that about 60% of amino acids in VASP are disordered (5), we expect VASP to be a flexible crosslinker that can bind over long distances. Therefore, we model VASP as a spherical crosslinker of radius 30 nm with four F-actin binding sites distributed across the surface of the sphere (Figure 1B). Cytosim also requires specification of a binding distance parameter between VASP and actin. This binding distance parameter controls F-actin binding propensity – the reaction propensity is calculated based on the abundance of F-actin found within the sphere centered on the binding site with radius determined by binding distance. To determine the appropriate binding distance, actin networks were simulated at various crosslinker binding and unbinding rates at binding distances 5, 10, 15, 20, and 30 nm respectively. We then chose a binding distance of 30 nm because this value allowed us to obtain all the experimentally observed actin networks.

#### Position evolution

CytoSim employs the Langevin equation to evolve the position of points (both along the fiber and those of VASP tetramers) considered. The discretized points along the filament and the VASP tetramers are considered. For a system of N particles, there are 3N coordinates. For a particle i, the coordinates are given by x_i_ = {x_i1_, x_i2_, x_i3_}. The position along each of the dimensions j is evolved according to the stochastic differential equation given by,

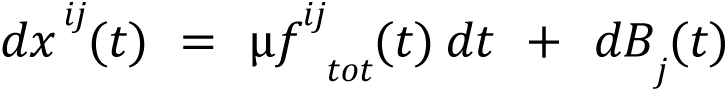

Here, μ is the viscosity of solvent, and *f*^*ij*^_*tot*_ (*t*) represents the total force acting on the particle at time t. The diffusion term (noise) is given by a random variable sampled from a distribution with mean 0 and standard deviation 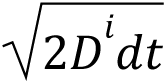, where the diffusion coefficient is given by µ*k*_B_*T* where T is temperature and k_B_ is Boltzmann constant. Please refer to Supplementary Table 1 for a detailed description of parameters used in this study.

#### Steric considerations

While the high-density increases propensity of chemical reactions (actin-VASP crosslinking), the steric-driven reduction in mobility within the droplet medium can either increase or decrease the effective rate constant (74). To capture this effect, we employ a steric repulsion potential between VASP molecules, and actin filaments.

#### Actin-covered surface fraction of the droplet

An icosphere corresponding to the radius of the droplet is generated. The icosphere is sub-divided 4x to approximate the curvature of the spherical surface. This was done so the triangle side is comparable to the segmentation length of actin filaments (100 nm). Actin filaments were discretized to the level of monomers and the monomers within 100 nm of the subdivided icosphere were considered to be close to the surface. Each of the monomers were assigned to the icosphere surface triangle closest to it. Thus, every snapshot from the Cytosim simulations was converted to a surface density plot and the fraction of occupied triangles is reported as the actin-covered surface fraction over time.

#### Simulations studying the role of droplet deformation

To understand the role of actin cytoskeleton in droplet deformation, we build on our previous hypothesis that the droplet deformation is driven by imbalance between droplet surface energy and actin ring bending energy. Therefore, in this study, we simulate the droplet deformation using the following protocol. The actin filaments are simulated for 600s under elongation rates and VASP-actin kinetics that favor ring formation. Once the maximum filament length is achieved, the network no longer elongates. Subsequently, we alter the droplet shape iteratively by increasing the Z-axis span of the volume in steps of 10 nm under constant volume conditions. The ring and diffusing VASP tetramers are allowed to equilibrate within the deformed spheroidal boundary shape for 1s (in Langevin time steps of 2ms) before undergoing subsequent deformation. The equilibration is achieved by a combination of diffusive dissipation through Langevin dynamics and chemical dissipation from VASP-actin (un)binding resulting in bundle reorganization. As the largest filament length studied is 2π microns, the highest aspect ratio a/b is achieved when 2a=2πRdrop (when R=1 mum, (a/b)max∼5.56). The droplets are elongated till we achieve (a/b)max

#### Determination of time between two deformations

To determine the time between two 10nm deformations, we deformed VASP-only droplets at various rates. At faster rates, we observe that the VASP-droplets do not equilibrate by diffusion adequately and are characterized by a region of the droplet without any VASP tetramers. Thus, our droplet deformation approach is a quasi-static approximation of the non-equilibrium process observed in experiments.

#### Determining maximum aspect ratio

Our simulation procedure approximates the droplet deformation process that is driven by competition between actin ring bending forces and the droplet surface tension. Thus, contact of actin along the droplet boundary, particularly the high-curvature caps (ends of the major axis) of the spheroidal droplet is critical. Thus, we need to determine a threshold volume fraction to identify the high-curvature caps and also choose an actin concentration threshold. In this study, we empirically choose a volume fraction of 12.5% and actin concentration threshold to be C_bulk_/3. The droplet is deemed to have reached the maximum aspect ratio when the actin concentration within the spheroidal caps falls below the concentration threshold.

**Figure S1.**
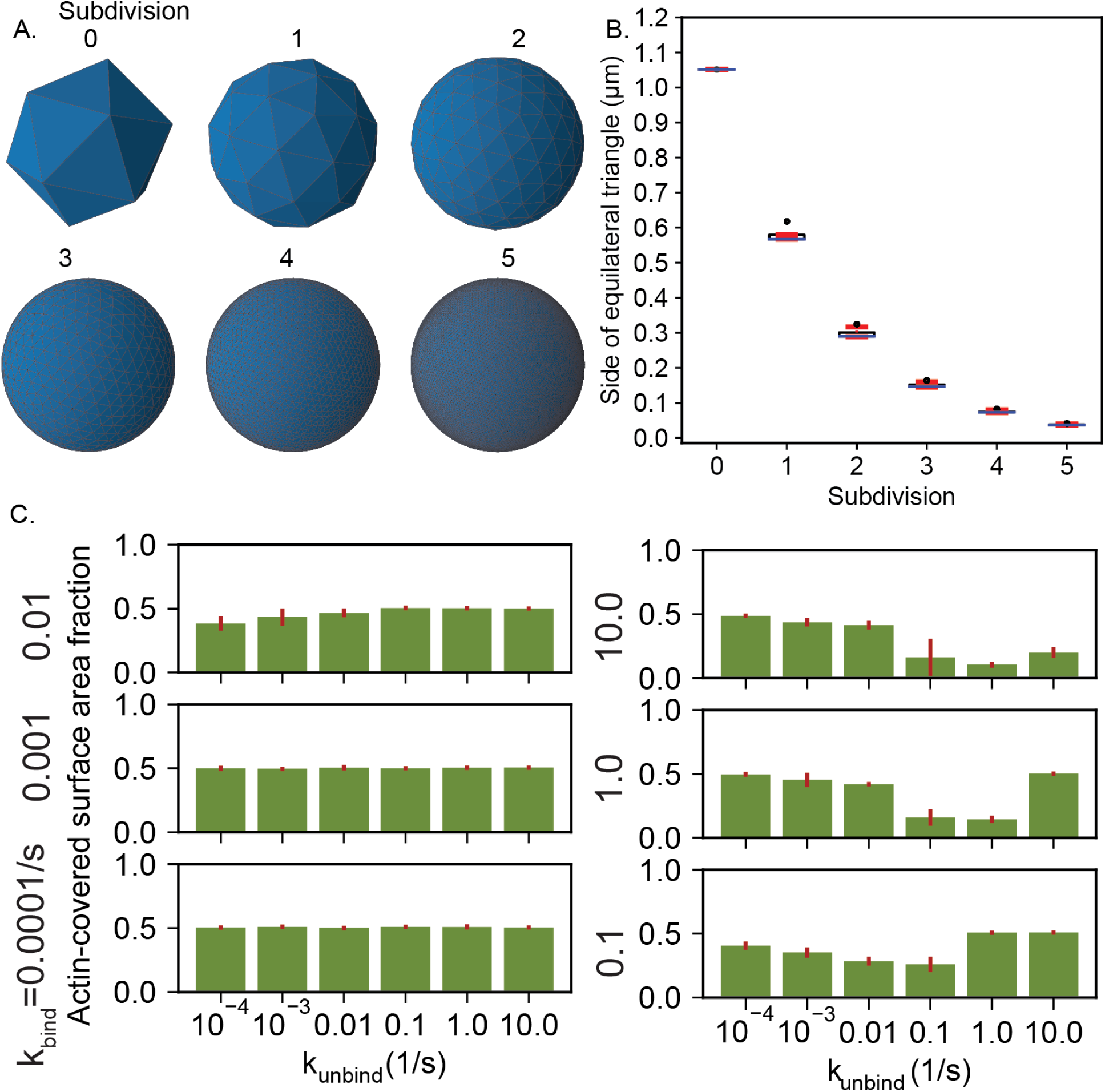
Steps used to obtain area fraction of actin. A. Icospheres generated at various subdivision values. B. Box plot showing the distribution of the triangle side as a function of subdivision of the icosphere. Median (blue), quartiles (black), 95% confidence range (red) and outliers (black circles) are shown for each level of subdivision. C. Actin-covered surface area fraction is calculated by generating a surface actin density map as described in Supplementary Methods and shown in Figure 3. Bar graphs mean and error bars correspond to standard deviation of final actin-covered surface fraction under various unbinding and binding rates. Data from the last 30 frames from each of the three replicates were used.

**Figure S2.**
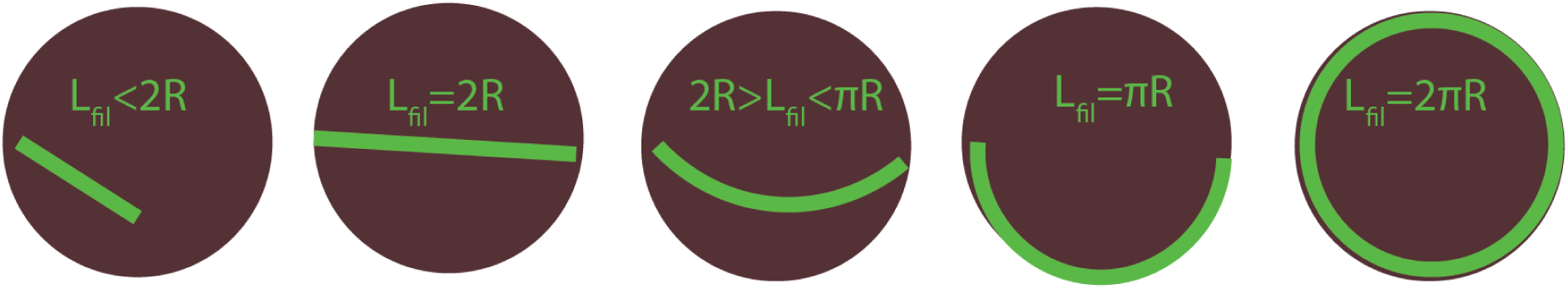
Schematic description of the minimum-energy configurations within spherical droplet for actin filaments of various lengths.

**Figure S3:**
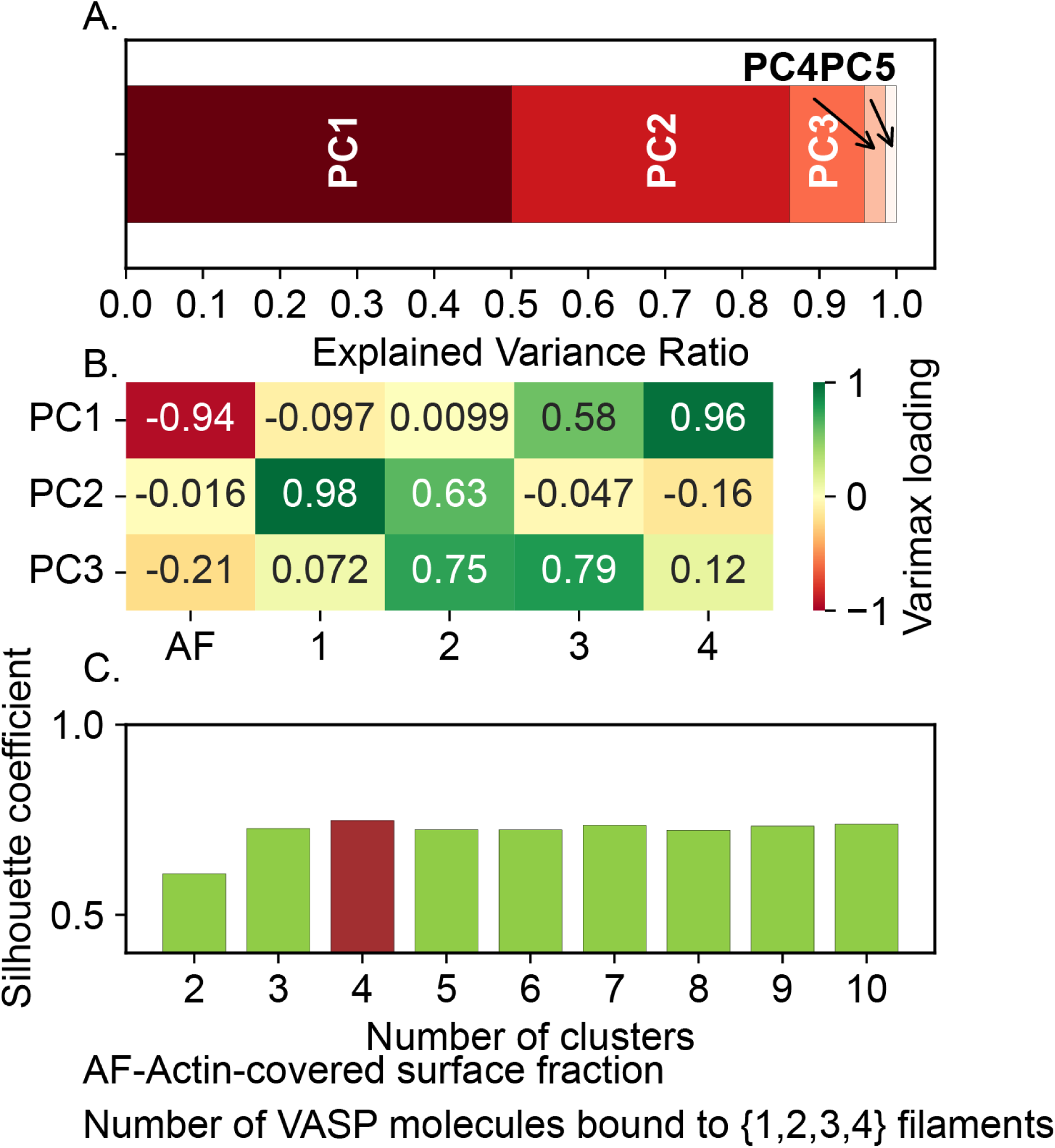
Feature and cluster number optimization to determine salient actin shapes present in our simulation. A. Principal component analysis on the five order parameters shows that the first two PCs explain 95.83% of variance. B. Varimax loading of the first two PCs suggests that the first PC takes information of Actin-covered surface fraction, and fraction of VASP molecules bound to {3 and 4} filaments while the second PC is dominated by information from the fraction of VASP molecules bound to {1, 2} filaments respectively. C. Silhouette coefficient was calculated to find the optimal number of clusters in our dataset.

**Figure S4.**
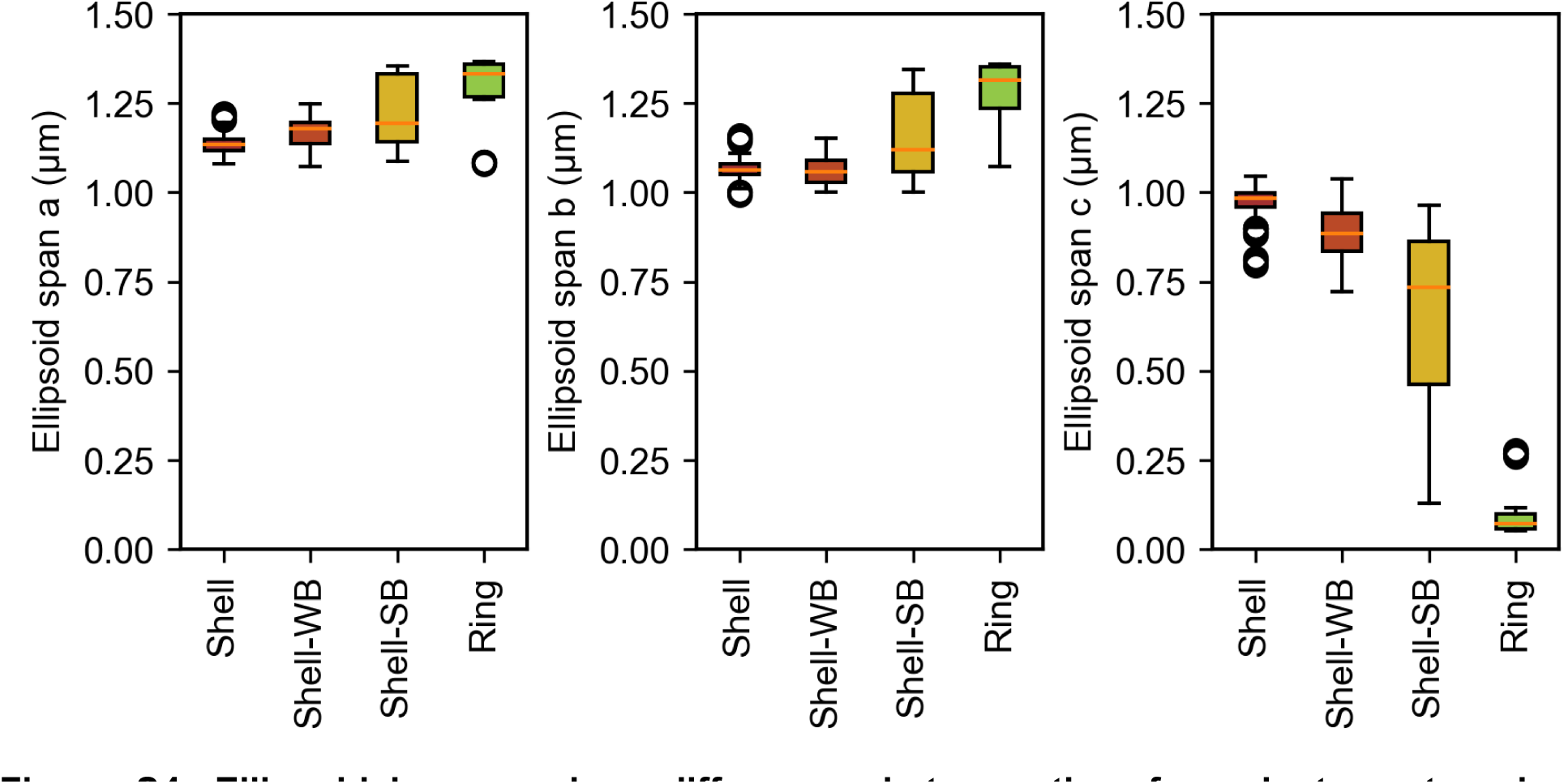
Ellipsoidal spans show differences between the four cluster categories. Assuming the actin network shape to be ellipsoidal, the dataset shown in Figure 3E was analyzed. The resulting spans (a≥b≥c) are shown here as boxplots. The cluster category is mentioned along the X-axis (Shell-WB-Shell with weak bundling, Shell-SB - Shell with strong bundling). Median (orange line), quartiles (top, bottom edges of the box), 95% confidence interval (whiskers) and outliers (open circles) are shown.

**Figure S5.**
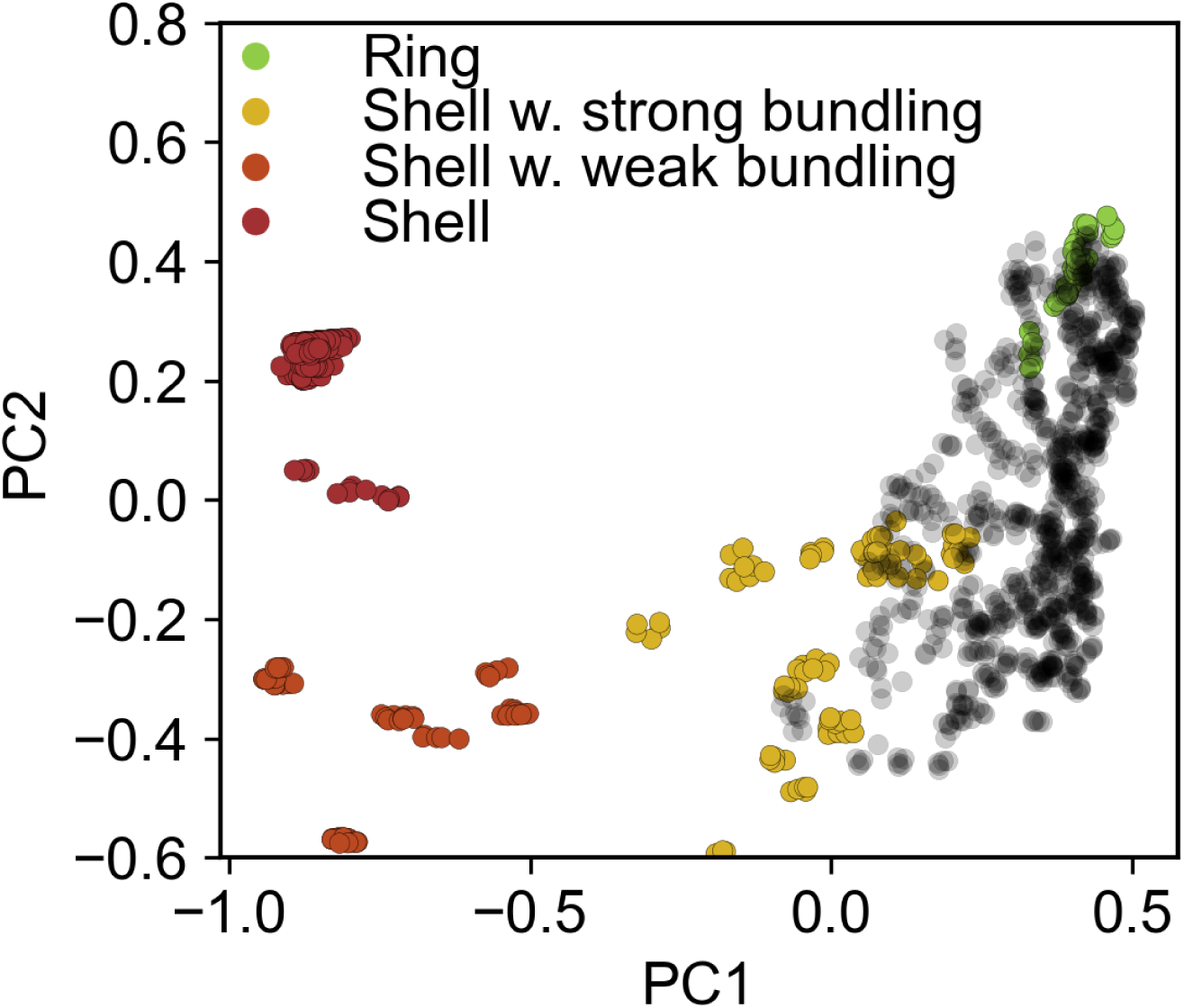
Data overlay highlights relevant shapes sampled by simulations at various T_2πR_. Data from simulations at various k_bind_ and k_unbind_ shown in Figure 2 (dataset 1) are combined with data from simulations at various T2πR shown in Figure 5 (dataset 2). The first two principal components are plotted. Data from dataset 1 are colored by their cluster identity while the data from dataset 2 are shown as black circles.

**Figure S6.**
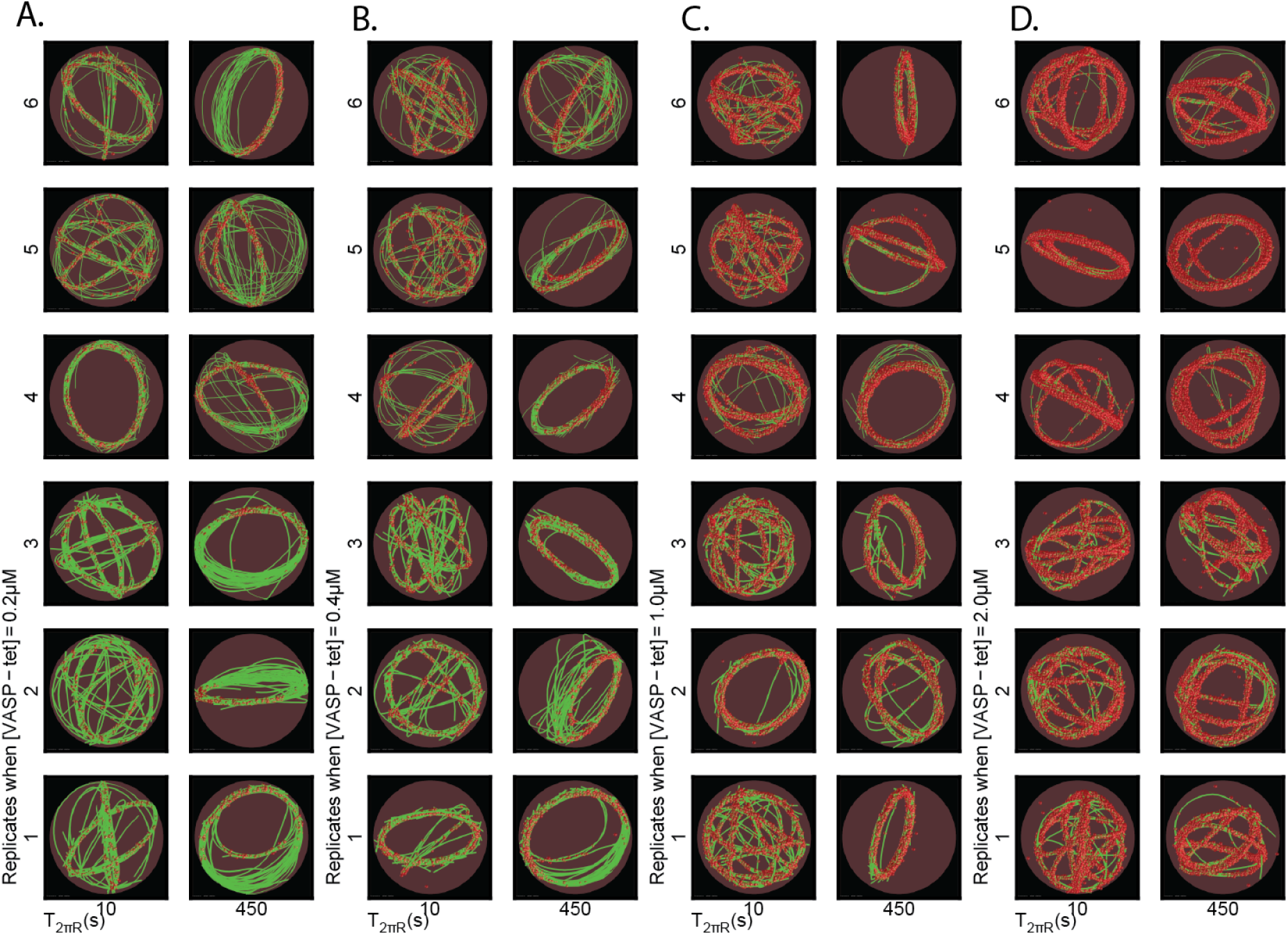
Heterogeneity in final network shapes at T_2πR_=10s and 450s shown for tetramer concentrations A. 0.2 μM, B. 0.4 μM, C. 1.0 μM and D. 2.0 μM (5 replicates). Actin filaments are shown in green while VASP tetramers are shown as red spheres. R_drop_ = 1 μm.

**Figure S7.**
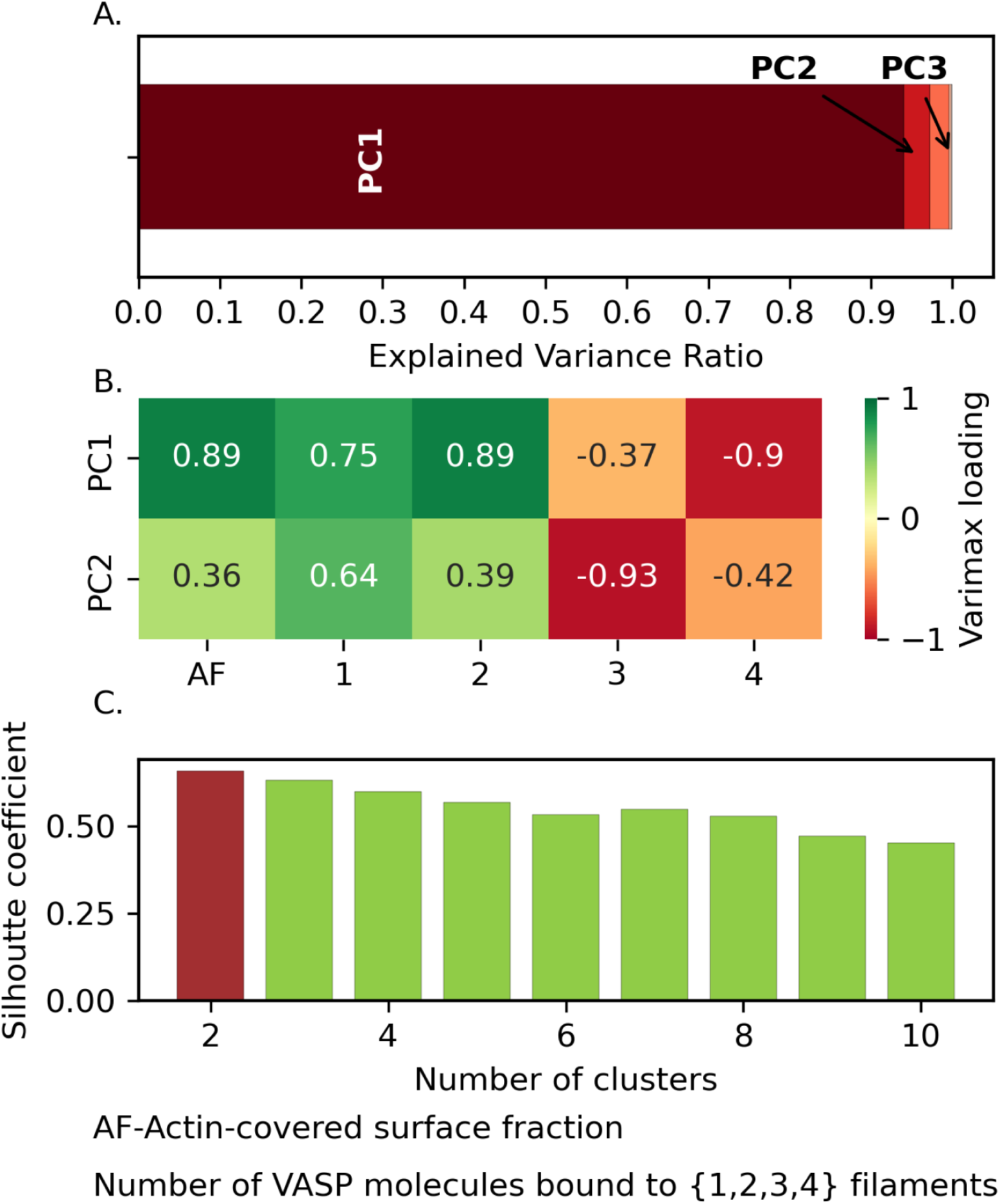
Feature and cluster number optimization to determine salient actin shapes present in our simulations at various T_2πR_. Data corresponding to rings, and rings with strong bundling from Figure 2 were combined with data from simulations at various T_2πR_ (Figure 5). A. Principal component analysis on the five order parameters shows that the first two PCs explain 99.06% of variance. B. Varimax loading of the first two PCs suggests that the first PC takes information of fraction of VASP molecules bound to {2,3, and 4} filaments while the second PC is dominated by information of Actin-covered surface fraction, respectively. C. Silhouette coefficient was calculated to find the optimal number of clusters in our dataset.

**Figure S8.**
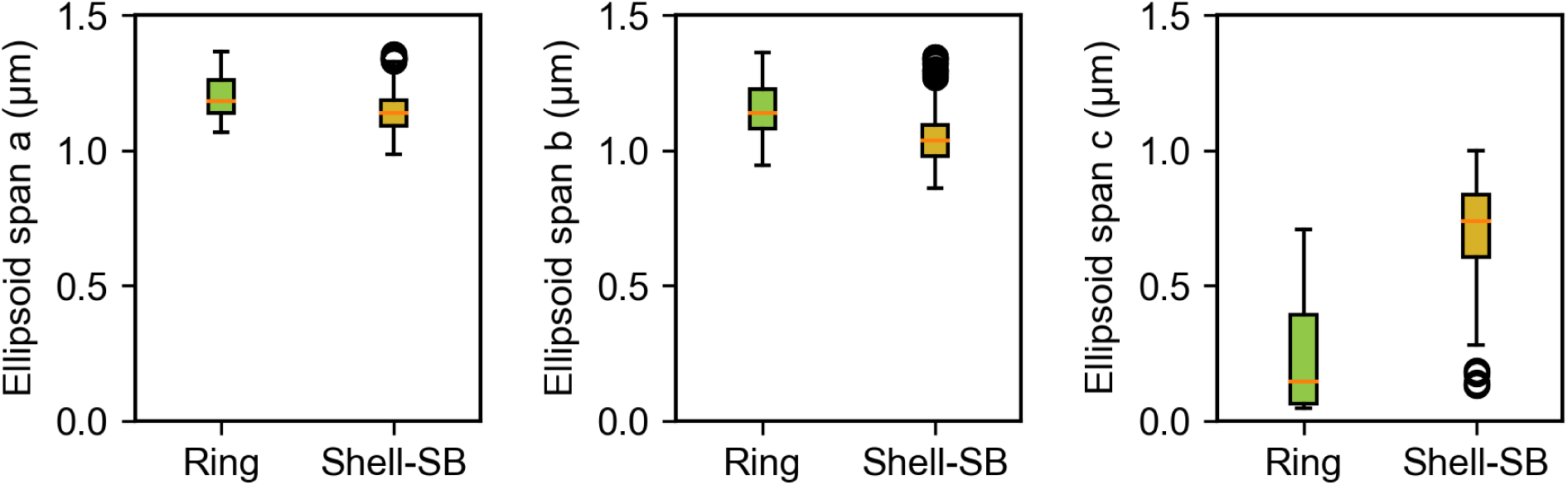
Ellipsoidal spans show differences between Ring and Shell-SB. Assuming the actin network shape to be ellipsoidal, the dataset shown in Figure 5C was analyzed. The resulting spans (a≥b≥c) are shown here as boxplots. The cluster category is mentioned along the X-axis. Median (orange line), quartiles (top, bottom edges of the box), 95% confidence interval (whiskers) and outliers (open circles) are shown.

**Figure S9.**
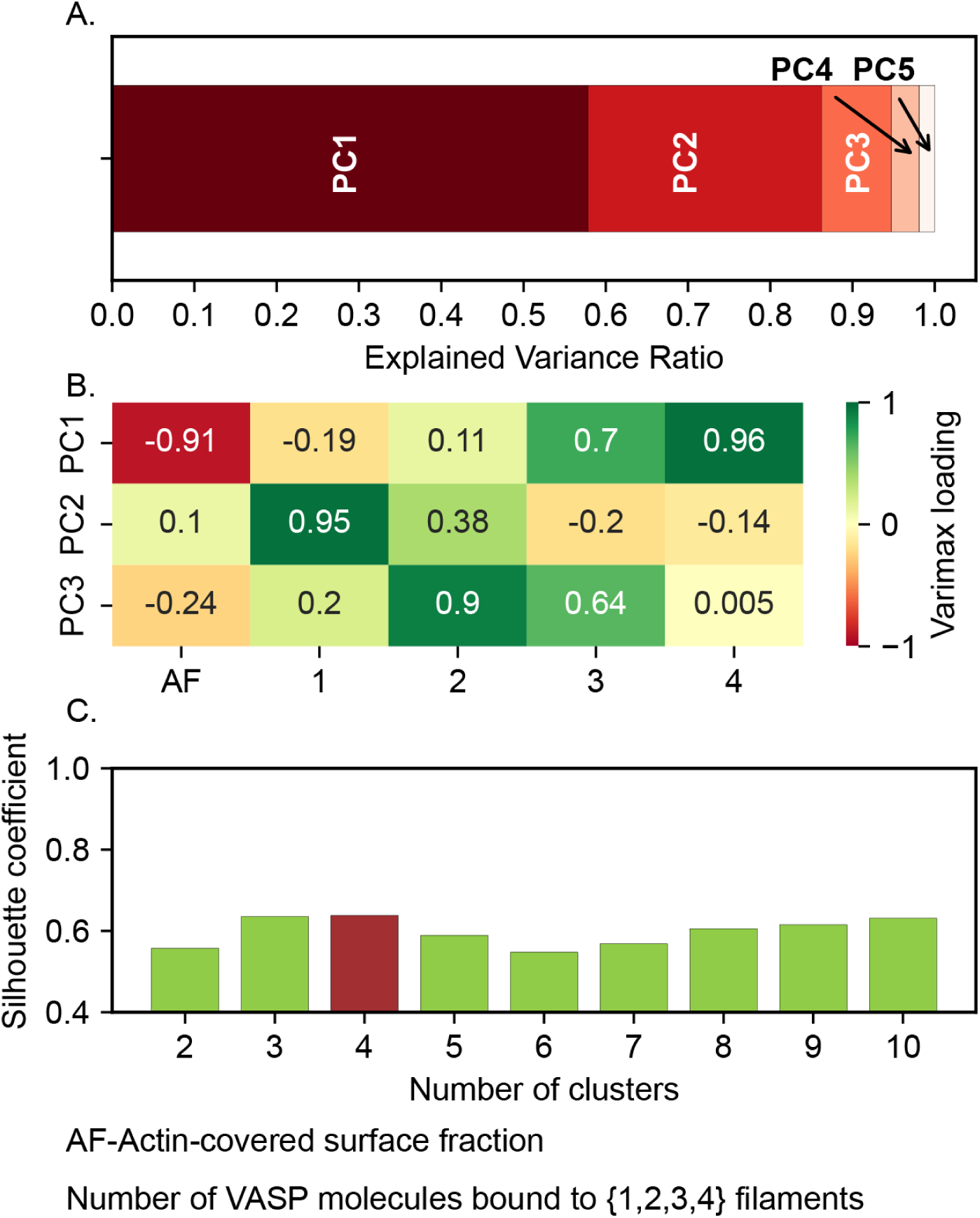
Feature and cluster number optimization to determine salient actin shapes present in our simulations with varying k_unbind_. A. Principal component analysis on the five order parameters shows that the first two PCs explain 94.72% of variance. B. Varimax loading of the first two PCs suggests that the first PC takes information of Actin-covered surface fraction, and fraction of VASP molecules bound to {3, and 4} filaments while the second PC is dominated by information from the fraction of VASP molecules bound to {1, and 2} filaments respectively. C. Silhouette coefficient was calculated to find the optimal number of clusters in our dataset.

**Figure S10.**
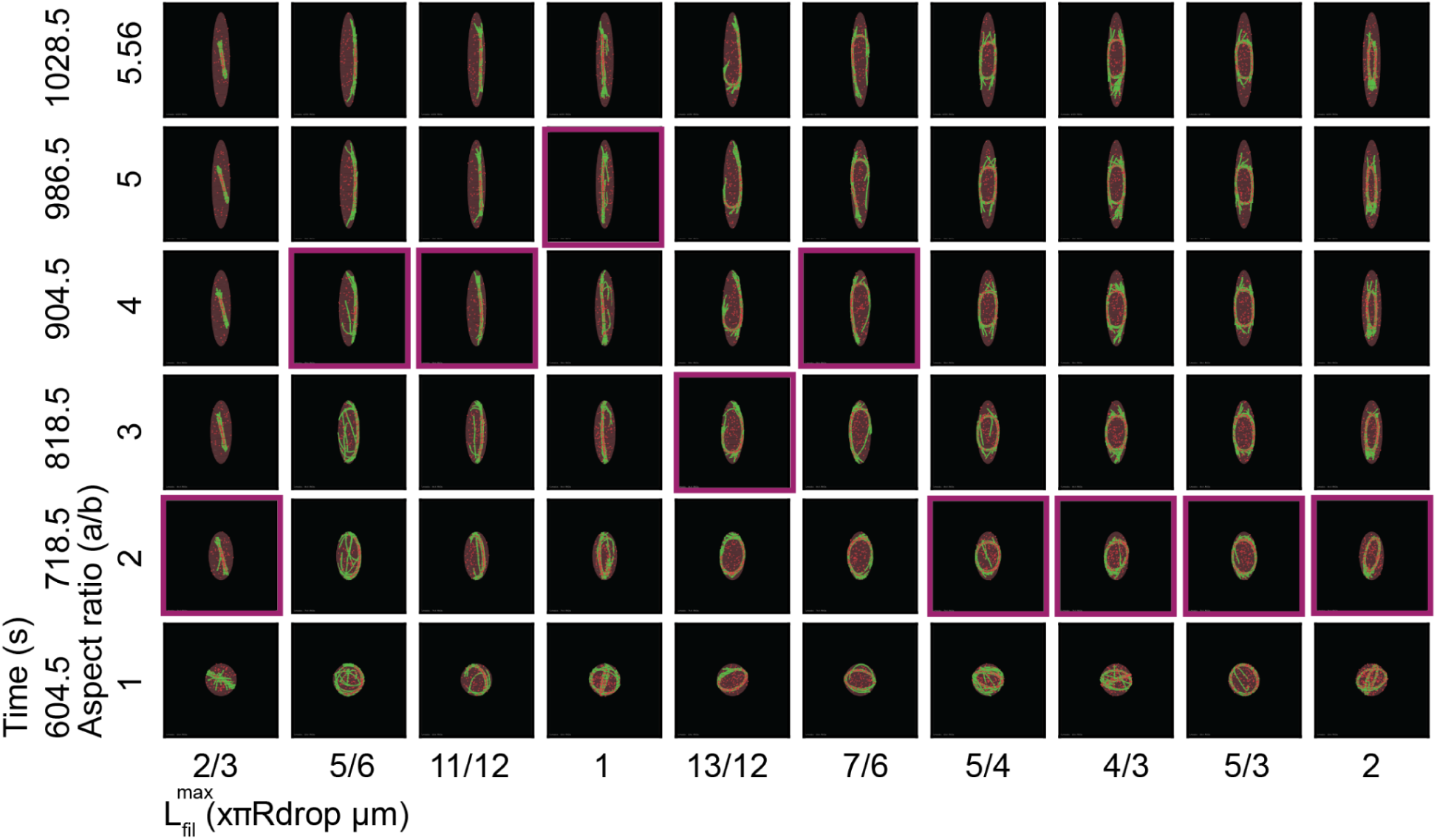
Snapshots from trajectories at various L_fil_^max^ are shown at specific time points along with the corresponding aspect ratios. Snapshots with integral aspect ratio closest to the maximum aspect ratio are colored in purple. Each subpanel shows actin filaments (green), VASP-tetramers (red spheres), within the droplet volume shown in brown. The maximum aspect ratio was determined as mentioned in Supplementary Methods.

**Figure S11.**
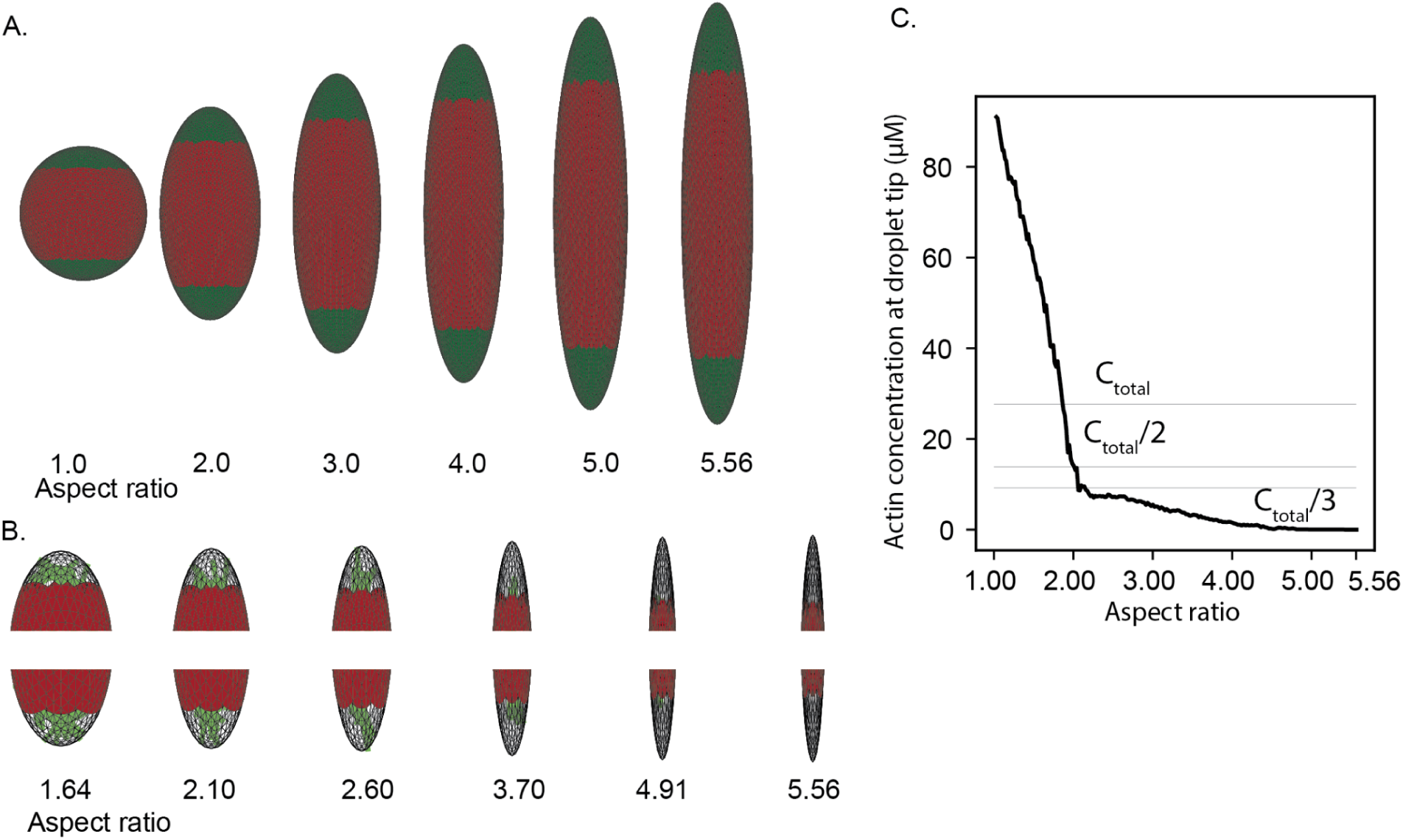
Determination of maximum aspect ratio. A. Droplets of various aspect ratios are shown. B. The top and bottom caps of the droplet are shown at various aspect ratios from simulations where the Lfilmax is π μm. 12.5% (6.25% in the top and 6.25% in the bottom) of the droplet volume is shown as transparent subvolume along with actin filaments (green). C. The total actin concentration within the caps spanning 12.5% subvolume is shown at various aspect ratios. Dotted lines represent 100%, 50%, and 33.34% of the total actin concentration within the droplet. The threshold of 33.34% and subvolume of 12.5% were chosen to identify the maximum aspect ratio of the deforming droplet. Above the maximum aspect ratio, we deem the deformation physically unrealistic.

### Movie Captions

Movie M1. Movie showing representative trajectories showing actin filaments (green) growing (growth rate = 10.3nm/s) within R_drop_=1um droplets of VASP-tetramers (red spheres) at various k_bind_ (mentioned on the left) and k_unbind_ (mentioned on the bottom) values. T_sim_=600s, Δt_frame_ = 5s, N_filaments_ = 30.

Movie M2. Movie showing representative trajectories of VASP tetramers (red spheres) simulated under the same actin addition rate but with different maximum filament lengths under ring forming conditions (k_bind_ = 10.0/s and k_unbind_ = 1.0/s) within R_drop_=1um droplets. T_sim_=600s, Δt_frame_ = 5s.

Movie M3. Gallery of representative trajectories showing actin filaments (green) growing at different filament extension rates specified by the time it takes for filaments to reach 2πR_drop_ (in seconds) at various [VASP-tet] concentrations within R_drop_=1um droplets. VASP tetramers are shown as red spheres. T_sim_=600s, Δt_frame_ = 5s, N_filaments_ = 30.

Movie M4. Gallery of representative trajectories showing VASP tetramers (red spheres) simulated under the same actin addition rate but with different maximum filament lengths under ring forming conditions (k_bind_ = 0.1/s and k_unbind_ = 0.1/s). Actin filaments were simulated for T=600s within a spherical boundary (Video shows 2 snapshots from this part of the simulation) Later, the aspect ratio of the droplet was changed by 10nm each step (at constant volume) and the system was allowed to adapt for 1s before further deformation. Video sampled at 1 frame per second.

## Notes

### Competing Interest Statement

The authors have declared no competing interest.

